# MonoAlg3D: Enabling Cardiac Electrophysiology Digital Twins with an Efficient Open Source Scalable Solver on GPU Clusters

**DOI:** 10.1101/2025.04.09.647733

**Authors:** Lucas Arantes Berg, Rafael Sachetto Oliveira, Julia Camps, Zhinuo Jenny Wang, Ruben Doste, Alfonso Bueno-Orovio, Rodrigo Weber dos Santos, Blanca Rodriguez

## Abstract

Modelling and simulation are essential in biomedicine, and specifically in computational cardiology. Reliable, efficient and accurate solvers are critical. This study presents an open source, GPU-based cardiac electrophysiology solver for scalable digital twin multiscale simulations (MonoAlg3D), incorporating conduction system calibration and performance optimization. The solver employs the monodomain equation coupled with the Purkinje network, solved via the finite volume method, featuring a GPU-based linear solver and concurrent simulation dispatch with MPI. We demonstrate a 10.94× speedup over a CPU-based solution and scalability by running 512 simulations on 128 compute nodes, completing all coarse-mesh simulations in less than 24 minutes and fine-mesh simulations in 303 minutes. We also demonstrate integration into a cardiac digital twin pipeline for personalisation based on clinical data. The proposed open source solver enhances computational efficiency and physiological fidelity, enabling large-scale, high-speed cardiac simulations. This work marks a significant step toward fast and scalable cardiac simulations on GPU architectures, with integration in a Digital Twin personalisation pipeline including the conduction system.

## Introduction

Computational modelling and simulation techniques in biomedicine have advanced over the last decades, from enabling investigations of disease mechanisms to making in silico trials for therapy evaluation possible^1−3^. Computational cardiology is a field that exemplifies a substantial amount of progress ^4−7^. Credible computer models of the heart are now available from subcellular to whole-organ dynamics, and efficient and accurate solvers have also been made available enabling large simulation studies. These two advances have driven the field forward towards the realisation of the the ‘Digital Twin’ vision in healthcare ^7−10^ and in silico trials for therapy evaluation^11^. For this, cardiac models need to incorporate clinically-relevant features of cardiac function and structure, such as the Purkinje conduction system, fibre orientation, anisotropy, cell coupling, and ECG computation, among others. Moreover, *in silico* clinical trials and therapy evaluation require consideration of large cohorts of virtual patients. Given ecological and economical limitations in computational resources, simulation software needs to provide an accurate and efficient approximation to the mathematical models describing such phenomena. Finally for reproducibility purposes, the software and models need to be made open source.

The challenge of the high computational costs associated with the resolution of cardiac models has led to the development of more efficient numerical schemes and the adoption of parallel computing techniques to reduce simulation times. In addition, the inclusion of the Purkinje system is a fundamental step toward physiological accuracy and correct modelling of ventricular activation in cardiac digital twin models. In experimental and computational studies, it has been shown that this structure can initiate and maintain certain types of arrhythmias due to altered conduction properties under pathological conditions leading to ectopic beats and reentrant circuits^12^.

Equally important, open source software presents several advantages ^13^: transparency (as the source code is freely available to the public, allowing end-users to validate and verify its functionalities), as well as flexibility and modularity (users can adapt and customise the software to their particular needs by adding novel features or implementing new modules that can be shared in a collaborative development environment). Along these lines, the cardiac modelling and simulation community has contributed several solvers to address these challenges^14−16^. OpenCARP^17^ and *li f e*^*x*^-*ep*^18^ constitute instances of more recent open source cardiac solvers based on central processing units (CPUs). However, despite providing substantial functionality, features such as the simulation of the Purkinje system remain to be addressed.

This study presents key improvements to MonoAlg3D, an open source, GPU-based solver for cardiac electrophysiology simulations, based on ^19^. These enhancements significantly improved computational performance and broaden its applicability. The widespread availability of Graphical Processing Units (GPUs) enables substantial speed-ups, which are essential for large-scale in silico trials ^20−24^. In addition, GPU-based solvers offer a cost-effective alternative by reducing reliance on extensive CPU resources in high-performance computing (HPC) environments. With the increasing adoption of GPU clusters, driven in part by advancements in artificial intelligence, these improvements further support efficient large-scale cardiac simulations.

MonoAlg3D uses the finite volume method (FVM) to simulate the monodomain model on GPU and/or CPU hardware, and OpenMP and NVIDIA CUDA to respectively parallelise CPU/GPU computations. Advancements include, firstly, a fully integrated Purkinje network model that allows for retrograde propagation, enabling investigations on its potential role in promoting and sustaining complex arrhythmias. Secondly, a novel solver for the diffusion linear system is implemented directly on GPUs to reduce the computational time of this component. Thirdly, a new output format is included to optimise disk space and access. Finally, the framework is expanded to support parallel dispatching across exascale HPC infrastructures, thereby facilitating large-scale, high-throughput simulations studies essential for digital twin and personalised medicine applications. Performance improvements are evaluated on different hybrid CPU/GPU combinations for the device and on two human-based cell models of increasing complexity, as well as compared to space adaptivity features presented in previous work^19^. Finally, to demonstrate the full capabilities of the proposed solver and verify its scalability on GPU clusters under more realistic scenarios, our last experiment presents a cardiac digital twin application considering a biventricular simulation with the Purkinje system and ECG recordings.

## Methods

### Monodomain model

The monodomain model is commonly used to describe electrical propagation due to its lower computational cost compared to the bidomain model^22,25^. In the next equations we present the mathematical models for the myocardium and the Purkinje system, along with their coupling. We use subscripts *P* for the Purkinje domain, *M* for the myocardium domain, and *d* ∈{*P, M*} for the full domain.

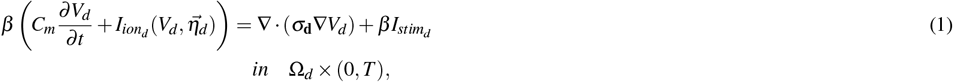

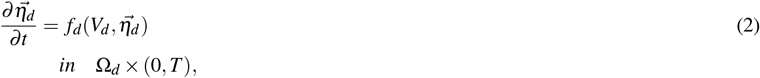

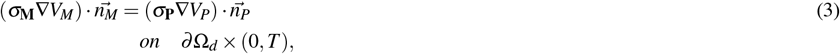

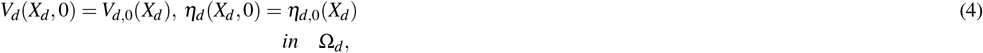

where *V*_*d*_ is the transmembrane potential of either domain, 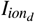 the total ionic current associated to the cellular model that depends on state variables 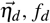 the non-linear system of equations encapsulating the dynamics of the state variables, *β* the surface-to-volume ratio, *C*_*m*_ the membrane capacitance, *σ*_**d**_ the domain conductivity tensor, and 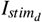 an external stimulus. The model is further closed with appropriate Neumann boundary conditions to ensure flux continuity between the myocardium and Purkinje domains as given by equation (3), where 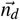 is the normal vector of the myocardium or Purkinje domain surfaces, *∂* Ω_*d*_. For myocardial surface nodes not coupled to the Purkinje system, equation (3) simply reduces to a standard non-flux boundary condition. Initial conditions are provided by equation (4). For the Purkinje domain we consider the one-dimensional form of equation (1), while for the myocardium domain its three-dimensional formulation.

Cardiac tissue is known to be comprised of strongly coupled fibres with anisotropic conduction properties. Such fibres are defined for each myocardial element by three orthonormal vectors 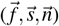, where 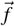 lies on the local fibre or longitudinal direction, 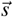 on the sheet or transversal direction, and 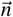 on the normal direction to the fibre. Moreover, associated with each of these vectors, there exist conductivity values *σ*_*f*_, *σ*_*t*_, and *σ*_*n*_, jointly defining the myocardial conductivity tensor as:

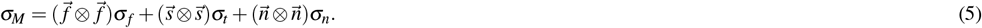

### Finite volume method applied to the monodomain model

A common technique to efficiently solve the monodomain model is to divide its reaction and diffusion parts using the Godunov operator splitting^26^. Applied to equations (1)−(2), this leads to the solution of two separate problems: a non-linear system of ordinary differential equations (ODEs)

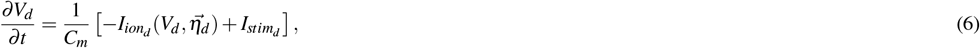

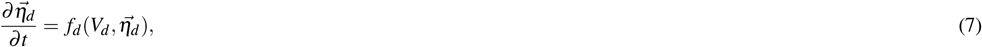

and a parabolic linear partial differential equation (PDE)

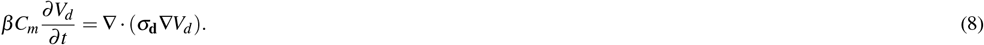

Within the different numerical techniques available, the FVM offers a robust approach for solving the monodomain model due to its foundation on conservative principles and applicability to diverse geometries^27^. This technique discretises the computational domain into control volumes. Each control volume is associated with a variable of interest, and the governing equations are applied to ensure the conservation of this variable across the control volume faces.

### Cell Model

For the solution of the cellular electrophysiology model described by Eqs. (6,7) MonoAlg3D offers different techniques for integration. It supports both the explicit Euler method as well as Rush-Larsen or other methods based on the generalization of matrix exponential, such as the Uniformization approach^28^.

### Myocardium modelling

To spatially discretise the diffusion term in equation (8), we consider the relation:

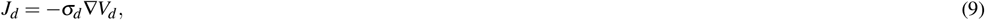

where *J*_*d*_ (*μA*/*cm*^2^) represents the density of intracellular current flow. Applying the divergence theorem and using equation (8), it yields:

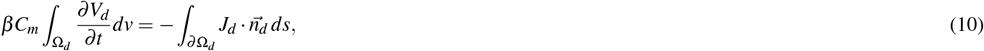

where 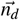 represents the normal vector to the domain surface. This equation is the basic term for deriving the linear system of equations associated with the linear PDE.

We now particularise the FVM equations for the myocardium. For simplicity, let us consider a tridimensional uniform mesh, consisting of hexahedra with a space discretisation *h*_*M*_. Located at the centre of each myocardial volume (*i, j, k*) is a node with the transmembrane potential *V*_*M*_ as the associated variable of interest. Assuming that the volumetric membrane current represents an averaged value in each hexahedron, and using (10), we then have:

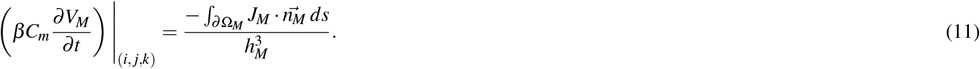

To support spatially varying fibre orientation and anisotropy of the myocardial conductivity tensor, the surface integral calculations in equation (11) consider the total sum of flows on the 6 faces of the control volume (each with face area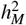) over a 27-neighbours stencil. This gives:

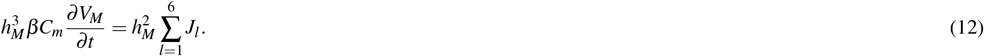

Each *J*_*l*_ in equation (12) is implicitly calculated by evaluating the spatial derivatives of *V*_*M*_ via second-order finite differences at timestep *n* + 1, and computing the average conductivity tensor given by (5) at the surfaces of the discretised volume. Altogether, the previous steps lead to a linear system to solve the diffusion equation using the backward Euler method. Additional details are provided in Supplementary Material section A.1.

### Purkinje modelling

To model the Purkinje system, we consider the one-dimensional form of the linear PDE given by equation (8), with time and space discretisations following an equivalent approach to the myocardial case presented above.

However, in the case of a Purkinje control volume, we have to consider three different possible configurations in our Purkinje networks (normal, branching, or terminal) as shown in Figure 1. The total flux given by equation (10) is:

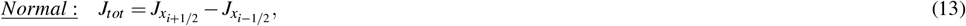

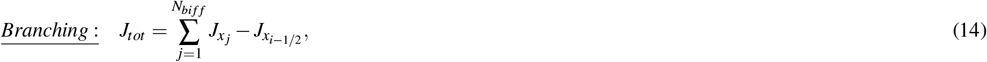

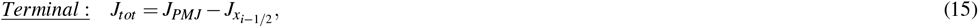

where *N*_*bi f f*_ is the number of Purkinje control volumes linked to the bifurcation, and *J*_*PMJ*_ is the flux associated at the Purkinje-muscle-junctions. Following a similar approach to the three-dimensional case, the fluxes 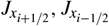, and 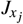 are calculated via finite differences at timestep *n* + 1 alongside the Purkinje conductivity *σ*_*P*_ associated to the surface of the discretised Purkinje control volume using harmonic means.

**Figure 1.**
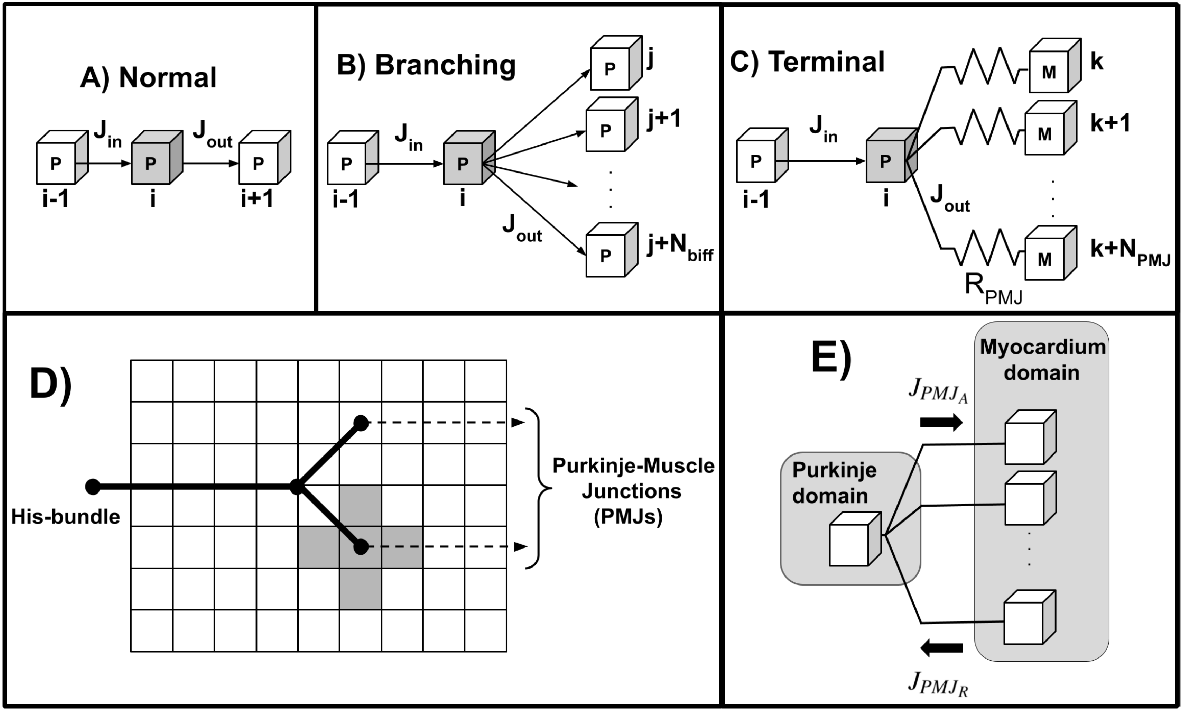
Illustration of the three possible configurations for a Purkinje control volume and the Purkinje coupling model. In panels (A)-(C), the control volume for which we are calculating the fluxes is depicted in grey. (A) Normal case, where a Purkinje control volume is associated with no branch. (B) Branching case, where a Purkinje control volume is linked to *N*_*bi f f*_ other Purkinje control volumes. (C) Terminal case, where a Purkinje control volume is coupled to *N*_*PMJ*_ myocardium control volumes from the myocardium domain by a fixed resistance *R*_*PMJ*_. Labels *P* and *M* indicate the domain where each control volume is located. (D) Simple Purkinje network with single bifurcation, coupling a Purkinje terminal to its five closest myocardium control volumes (coloured in grey). (E) Direction of the flux *J*_*PMJ*_ for anterograde 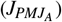 and retrograde 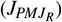 propagation, respectively.

### Purkinje−myocardium coupling

To model the coupling between Purkinje−myocardium domains, we consider an additional flux *J*_*PMJ*_. The electrical stimulus coming from the Purkinje system reaches the myocardium at specialised sites called Purkinje-muscle junctions (PMJs), spreading in the endocardium by a distance of approximately 1 *mm* between each other^29^. Importantly, PMJs are known to exhibit a characteristic asymmetric conduction delay due to electrotonic interactions of around 4 − 14 *ms* on the anterograde direction (Purkinje-to-myocardium)^30^ , and of about 2− 4 *ms* when propagation occurs in the retrograde direction (myocardium-to-Purkinje)^30^. This behaviour is characterised as a source-sink mismatch phenomenon since a single Purkinje terminal may need to activate a bulk of myocardium tissue^31,32^.

Typically, the PMJ coupling is modelled by a fixed resistance, linking a Purkinje element to several myocardium elements^33^. We follow this approach by modelling the flux *J*_*PMJ*_ (*μA*/*cm*^2^) using a fixed resistance *R*_*PMJ*_ and by coupling a single Purkinje control volume to its *N*_*PMJ*_ closest myocardium control volumes, as shown in Figure 1D. Moreover, the PMJ flux is given as a non-homogeneous Neumann boundary condition by:

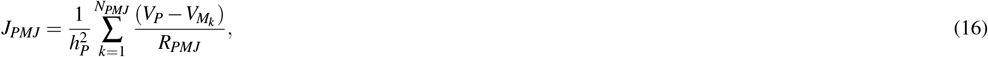

where the sign of the flux determines if *J*_*PMJ*_ exerts its action in the anterograde or retrograde direction (see Figure 1E).

### Numerical scheme

For the iterative solution of the coupled model, we start by solving the reaction terms describing the Purkinje and myocardium cellular models, given by the non-linear systems of ODEs in equations (6)-(7). We consider here the forward Euler method for simplicity, albeit MonoAlg3D is equipped with more advanced ODE schemes (such as Rush-Larsen and adaptive forward Euler). This gives:

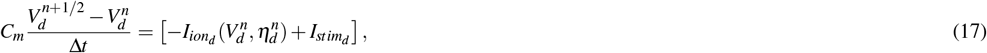

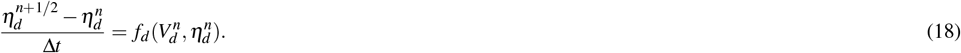

PMJ fluxes are computed next based on equation (16), as:

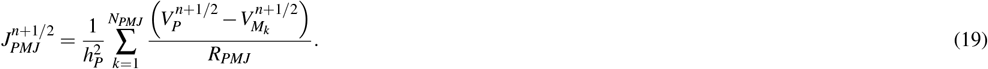

The diffusion terms of the Purkinje and myocardium domains involves the solution of the linear system:

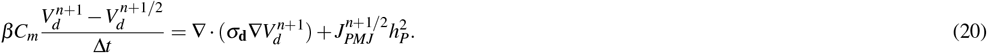

Observe that by computing *J*_*PMJ*_ at time *n* + 1/2, we decouple the Purkinje and Myocardium domains. This enhances the solver’s modularity, allowing different classes to be used for each domain, but at the cost of numerical stability. While a fully implicit solution of the PDE is unconditionally stable, decoupling the two domains results in a conditionally stable scheme, where *R*_*PMJ*_ constrains the maximum time step.

### ECG calculations

An approximation for the ECG can be computed by assuming that the tissue is immersed in an unbounded volume conductor^34^. The surface potential can be then calculated using the equation:

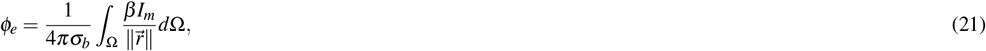

where *σ*_*b*_ is the bath conductivity, and 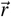 is the distance vector between source and field points, the latter essentially the electrode positions of the virtual ECG leads. The source term *β I*_*m*_ is given by the solution of the diffusive term ∇· (*σ* ∇*V*_*m*_), which is available in every timestep.

To efficiently implement this new functionality in MonoAlg3D, we implemented the calculations of the equation (21) using OpenMP in CPUs or CUDA on GPUs environments.

### Performance efficiency strategies

#### Solving diffusion on GPUs

In previous work^19^, the linear system linked to the diffusion term in equation (8) was exclusively solved in the CPU using an OpenMP version of the conjugate gradient (CG) method.To enable the solution of large linear systems on GPUs, we first converted its sparse matrix representation from the ALG format to a Compressed Sparse Row (CSR) data structure compatible with the cuSparse library. This allows to directly solve the CG on the GPU by using this data structure together with the methods implemented in the cuBLAS library. It is worth noting that the biconjugate gradient (BCG) method is also available in this new version.

#### New output format

To minimise disk space usage and improve output performance, we provide novel support for EnSight files as new output format. This format is also compatible with multiple visualisation tools, such as Paraview, allowing most post-processing workflows to be kept unchanged.

#### MPI batch

Finally, for sensitivity analysis and uncertainty quantification studies, MonoAlg3D provides a novel feature for the concurrent dispatch of multiple simulations using the message passing interface (MPI) standard. Given a baseline simulation and a range of parameters, the solver generates automated configuration files for all possible combinations of input parameters. Each configuration file is then dispatched in parallel using MPI. This allows to upscale more efficiently the number of jobs running in HPC environments, enabling for instance to perform hundreds of simultaneous simulations for a given patient using a wide a range of parameters. Such a feature is an important step towards *in silico* trials, drug therapy, and risk assessment studies.

### Computational simulations

Two sets of experiments were used to evaluate the improvements implemented in MonoAlg3D: a benchmark cuboid mesh to quantify performance improvements; a human biventricular mesh coupled to a Purkinje network (see Supplementary Figure S4A section A.3), as an exemplar of cardiac digital twin application.

All our numerical experiments were performed in the Polaris supercomputer provided by the Argonne Leadership Computing Facility, a 560 node HPE Apollo 6500 Gen 10+ system. Each computing node is equipped with a 2.8 GHz AMD EPYC Milan 7543P 32 core CPU with 512 GB of DDR4 RAM and four NVIDIA A100 GPUs.

#### Benchmark myocardium cuboid

A numerical test, adapted from Niederer et al.^35^, was conducted with minor domain size modifications to: 1) evaluate GPU speedups in solving non-linear ODEs and the parabolic PDE; 2) assess space adaptivity effects on efficiency; and 3) compare disk space usage between EnSight and VTK formats.

The test used a 1 × 1 × 1 *cm*^3^ myocardium cuboid with transverse anisotropic conduction *σ*_∥_ = 1.334 *mS*/*cm, σ*_⊥_ = 0.176 *mS*/*cm*), monodomain parameters *β* = 1400 *cm*^−1^, *C*_*m*_ = 1 *μF*/*cm*^2^, and human-based ventricular models: *ten Tusscher* (12 state variables)^36^ and *ToR-ORd* (43 state variables)^37^. Stimulation was applied in a 0.15 × 0.15 × 0.15 *cm* region for 2 *ms* at 53 *pA*/*pF*.

The nonlinear ODEs were solved using the Rush-Larsen scheme^38^ with Δ*t* = 0.01 *ms*, while the parabolic PDE used Δ*t* = 0.02 *ms* for a total of 1500 *ms*. Without space adaptivity, uniform discretization at *h*_*M*_ = 250 *μm* led to 64, 000 control volumes. Adaptive resolutions ranged from 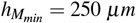 to 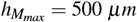, with refinement/de-refinement bounds at 10.01/10.00 and adaptation every 10 timesteps.

The benchmark was tested across six CPU/GPU configurations:

- <u>A+OC+PC:</u> Adaptive, ODEs on CPU, PDE on CPU;
- <u>A+OG+PC:</u> Adaptive, ODEs on GPU, PDE on CPU;
- <u>OC+PC:</u> Non-adaptive, ODEs on CPU, PDE on CPU;
- <u>OC+PG:</u> Non-adaptive, ODEs on CPU, PDE on GPU;
- <u>OG+PC:</u> Non-adaptive, ODEs on GPU, PDE on CPU;
- <u>OG+PG:</u> Non-adaptive, ODEs on GPU, PDE on GPU.

Mesh geometry and transmembrane potential were saved every 100 timesteps. More details on the benchmark setup are provided in Supplementary Figure S3 section A.2.

#### Cardiac digital twin with Purkinje network

To demonstrate the full capabilities of the proposed GPU cardiac solver, we conducted a simulation study within a biventricular cardiac digital twin pipeline incorporating a Purkinje network. The study had three objectives: 1) evaluate solver performance in realistic scenarios; 2) calibrate Purkinje coupling parameters *R*_*PMJ*_ and *N*_*PMJ*_ to physiological anterograde PMJ delays; and verify solver scalability for concurrent GPU simulations.

We used a human biventricular mesh (76-year-old female, 87 *kg*, 107 *cm*^3^ volume) reconstructed from MRI^39^, previously applied in clinical ECG personalization^40−42^. Supplementary Figure S4 section A.4, presents further anatomical details, including its Purkinje network coupling (Supplementary Figure S4A), fiber orientation field (Supplementary Figure S4B), subendocardial Purkinje coupling layers (Supplementary Figure S4C), and *I*_*Ks*_ scaling factor map for T-wave personalization^41^ (Supplementary Figure S4D).

For cellular electrophysiology, we used the *ToR-ORd* human-based ventricular model^37^ with modifications for T-wave personalization^42^: 50% *I*_*Kr*_ scaling, 5× *I*_*Ks*_ scaling^43^, and reducing *τ* _*jca*_ from 75 to 60 *ms*. The Purkinje domain was modeled with the human-based Purkinje *Trovato* model^44^. Both ODE systems were solved via the Rush-Larsen scheme with Δ*t* = 0.01 *ms*. PDEs used the same discretization step for a total simulation time of 600 *ms*. The stimulus protocol consisted of a single pulse applied at the His bundle (*N*_*cells*_ = 25) with 40 *pA*/*pF* amplitude and 2 *ms* duration. For more details about the mesh configuration refer to Supplementary Material sections A.3 and A.4.

To evaluate performance, we tested two myocardium space discretizations: a *coarse* mesh (*h*_*M*_ = 500 *μm*) with 855, 670 control volumes and a *fine* mesh (*h*_*M*_ = 250 *μm*) with 6, 845, 360 volumes. The Purkinje domain used a fixed *h*_*P*_ = 250 *μm* with 7, 948 volumes. Solver scalability was assessed across three simulation setups:

- <u>1N1S:</u> 1 node, 1 simulation;
- <u>1N4S:</u> 1 node, 4 concurrent simulations;
- <u>128N512S:</u> 128 nodes, 512 concurrent simulations.

A large-scale simulation study calibrated *R*_*PMJ*_ and *N*_*PMJ*_ within a physiological range using MonoAlg3D’s MPI batch processing (see Supplementary Material section A.5). We ran 512 concurrent simulations, varying *R*_*PMJ*_ and *N*_*PMJ*_ based on an initial calibration (see Supplementary Material section A.6). For the *coarse* mesh, *R*_*PMJ*_ spanned [100, 1300] *k*Ω (32 values) and *N*_*PMJ*_ [15, 50] (16 values). For the *fine* mesh, *R*_*PMJ*_ ranged from [500, 2300] *k*Ω. ECG comparison with clinical data was performed using Pearson’s correlation coefficient across all 8 leads (I, II, V1−V6).

## Results and discussion

### Myocardium cuboid benchmark

An initial test was conducted using the OC+PC configuration, which uses entirely the CPU without space adaptivity to solve both the ODE and PDE systems, to evaluate the optimum number of OpenMP threads for the selected HPC facility. Five simulations were executed per number of threads, considering the human ventricular cellular *ToR-ORd* model(Fig. 2A). The best total execution times were found for an optimal number of 8 OpenMP threads, leading to a ≈ 6.66 × efficiency speed-up, and enabling benchmark execution times around 30 minutes.

**Figure 2.**
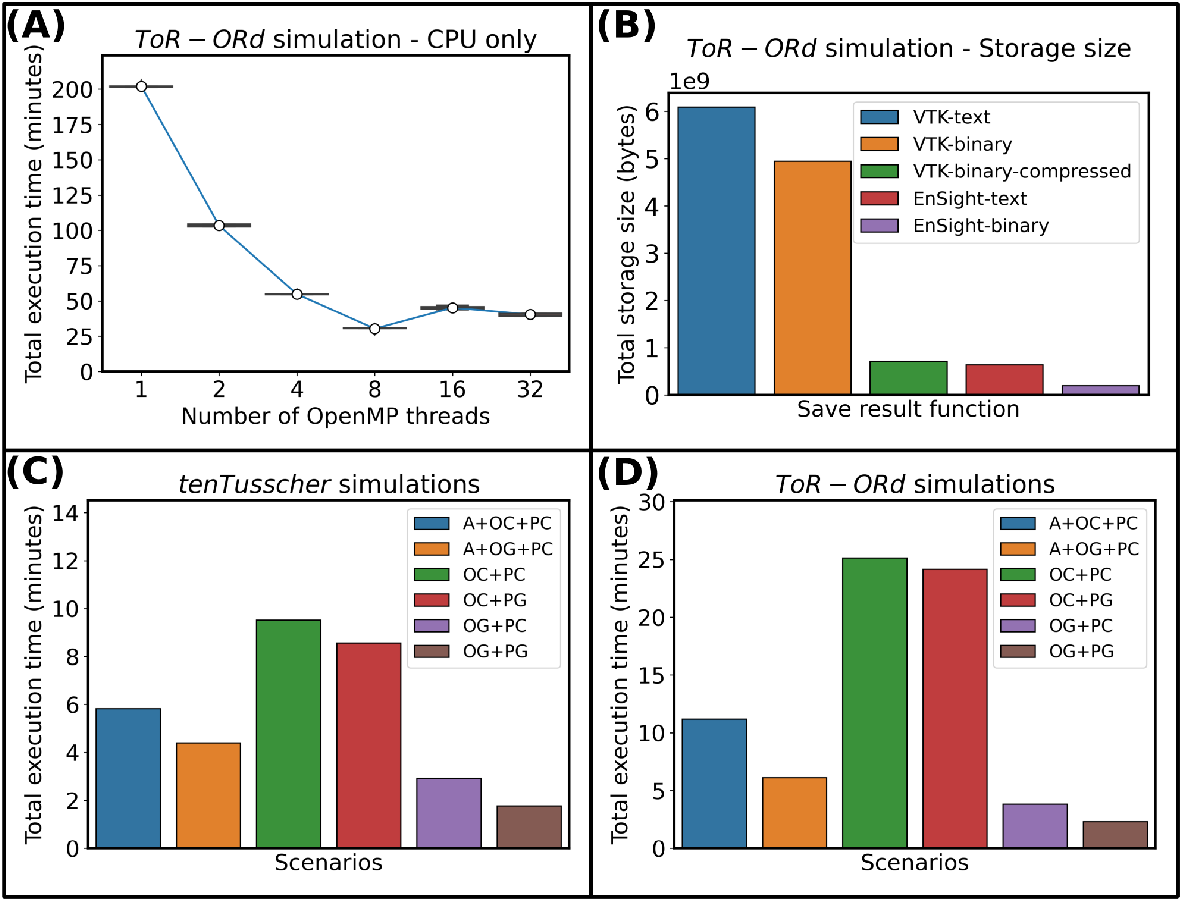
Results for the myocardium cuboid benchmark. (A) Execution time of the *ToR-ORd* simulation considering an OC+PC solver. (B) Disk usage for storing the mesh geometry and with the myocardial transmembrane potential for the *ToR-ORd* simulation using different file formats. (C) Total execution time for each of the 6 scenarios when using the *ten Tusscher* model. (D) Total execution time for each of the 6 scenarios with the *ToR-ORd* model.

Similarly, input/output efficiency was optimised by considering 5 broadly adopted scientific formats: VTK-text (ASCII), VTK-binary, VTK-binary-compressed, EnSight-text (ASCII), and EnSight-binary (Fig. 2B). The results from this analysis highlight substantial savings in output file size when saving each model state variable (transmembrane potential in our case) in EnSight-binary format. This resulted in file storage sizes of merely 0.19 *gigabytes*, while VTK-text required around 6 *gigabytes* to store the same outputs. Therefore, the EnSight-binary can save approximately 31× , 25× , 3.5× and 3.23× more disk space when compared to VTK-text, VTK-binary, VTK-binary-compressed, and EnSight-text formats, respectively.

We then evaluated the solver’s performance for each of the 6 considered combinations of CPU/GPU architectures (see section *Computational simulations/Benchmark myocardium cuboid*), using either the *ten Tusscher* (Fig. 2C) or the *ToR-ORd* (Fig. 2D) cellular models. Based on the analysis above, 8 OpenMP threads were used in all the cases. The efficiency results presented in Figure 2C for the *ten Tusscher* model yielded a maximum simulation time of ≈ 9 *min* for the OC+PC scenario (i.e., solving the entire problem in the CPU, without space adaptivity). Space adaptivity allowed the problem to be solved under 6 *min* in the CPU (A+OC+PC scenario), and in around 4 *min* if the ODE system was solved in the GPU (A+OG+PC scenario). The largest efficiency improvement was however found when the problem was solved entirely in the GPU (OG+PG scenario), decreasing the simulation time under 2 *min*. Equivalent results are presented in Figure 2D for the *ToR-ORd* model for all the CPU/GPU configurations. Solving the simulation entirely on the CPU without space adaptivity (OC+PC scenario) yielded the most demanding execution time of ≈ 25 *min*, while a 10.94× efficiency gain and a total simulation time below 3 *min* were attained by exploiting the full GPU implementation (OG+PG scenario).

The results above indicate that space adaptivity (scenarios A+OC+PC and A+OG+PG) did not lead to any improvements in performance when compared to solving the fully refined mesh entirely on the GPU (scenario OG+PG). This behaviour can be attributed to the computational overhead associated with spatial adaptivity, specifically reassembling the matrix of the PDE and updating the grid data structures. In contrast, preloading the fully refined mesh and transferring all the data structures for both the ODEs and PDE to the GPU at the onset of a simulation offers significant advantages in terms of memory usage and computational performance when a fixed spatial discretisation is used. This approach not only reduces data transfer between the host and the GPU, but also minimises memory allocation operations. Consequently, the computation at each time step is more regular than when space adaptivity is used, leading to an improved overall performance.

In addition, the joint analysis of the two considered cellular models revealed that solving the ODE system on the GPU was responsible for most of the performance gains. This becomes apparent by comparing scenarios OC+PC and OC+PG. For the *ten Tusscher* model, scenario OC+PG is just 1.11× faster than scenario OC+PC, while for *ToR-ORd* a speedup of 1.04× is obtained. However, when the ODE system was solved on the GPU (scenario OG+PC), a 3.26× gain was attained for the *ten Tusscher* model when compared to scenario OC+PC, while this improvement was even more pronounced for *ToR-ORd*, and around 6.60× gain. This result is primarily underlain by differences in algebraic complexity between both cellular models: the *ten Tusscher* model consists of 12 state variables, compared to 43 in *ToR-ORd*. Additionally, the *ToR-ORd* model involves a greater number of algebraic expressions, making it well-suited for GPU-based computations. Detailed execution times for both models can be found in the Supplementary Tables S1 and S2 section A.2.

### Cardiac digital twin with Purkinje network

Based on the results of our previous benchmark, all cardiac digital twin simulations were executed entirely on the GPU using 8 OpenMP threads, exploiting one GPU device for both the Purkinje and myocardium domains as this configuration demonstrated the best performance.

Figure 3 illustrates selected simulation results at *fine* mesh resolution for the identified optimal set of Purkinje coupling parameters from a total of 512 executions. This set was chosen based on similarity between the simulated and clinical ECG signals across all leads, as well as replicating a range of physiological anterograde PMJ delays across all Purkinje terminals when activating the myocardium. Nevertheless, the existence of other combinations of Purkinje coupling parameters yielding a similar activation pattern (Figs. 3A−3B) indicates that a range of cardiac digital twins can be approximated by the considered parameters, *R*_*PMJ*_ and *N*_*PMJ*_. Ventricular activation started ≈ 48 *ms* after His-bundle pacing (Figs. 3C−3D), with the whole biventricular domain being activated in around 110 *ms* (end of simulated QRS complex, Fig. 3E), also within the physiological range of 80 −120 *ms* for healthy subjects^29^. Moreover, the inclusion of heterogeneity in *I*_*Ks*_ from^42^ generated a reasonable approximation for the T-wave (Fig. 3E), with an average PCC of 0.81 across all the 8 independent ECG leads (I, II, V1−V6). The effects of anterograde PMJ delays were correctly recovered as shown in Figure 3D, as well as in Figure 3F for a PMJ site on the right ventricle. As it can be seen in the latter, the closest myocardial control volume to the Purkinje terminal could not be activated instantaneously, due to the source-sink mismatch between the Purkinje terminals and myocardial cells^31,32,45^. The myocardial control volume only became entirely depolarised approximately 4.34 *ms* after the stimulus reached the PMJ site, exhibiting an initial brief spike followed by a distinct blunted depolarisation (Fig. 3F). This behaviour, reported in different experimental studies^46,47^ and attributed to the electrotonic effects at the junctions, is further explored in the PMJ calibration results presented in Supplementary Material section A.6.

**Figure 3.**
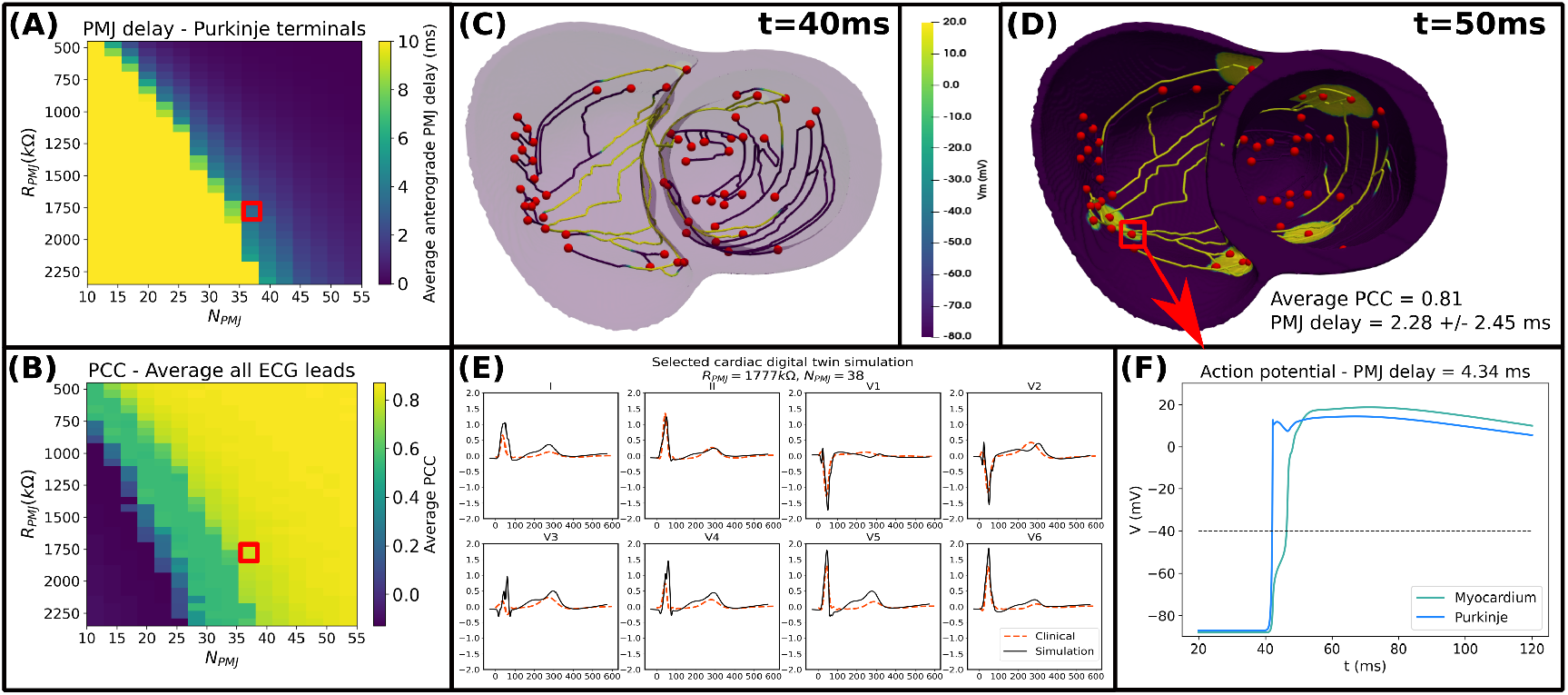
Results for the 512 digital twin simulations with the *fine* mesh. (A) Average anterograde PMJ delay across all Purkinje terminals. (B) Average PCC across all leads between the clinical and simulated ECG. Selected Purkinje coupling parameters (*R*_*PMJ*_ = 1777 *k*Ω, *N*_*PMJ*_ = 38) are highlighted by red squares. (C-D) Selected cardiac digital simulation with an average PCC of 0.81 in ECG reconstruction and average anterograde PMJ delay of 2.28 ± 2.45 *ms*, at times *t* = 40 and *t* = 50 *ms*, respectively. (E) Comparison between clinical and simulated ECGs. (F) Action potential upstrokes for the PMJ site highlighted in panel (D) with anterograde PMJ delay of 4.34 *ms*, at the terminal Purkinje volume (blue) and its closest coupled myocardium volume (lime).

In terms of scalability, Table 1 summarises execution times for our considered submission scenarios. For the 1N1S scenario (1 computing node, 1 simulation), total execution times for the *coarse* and *fine* discretisations were around 15 and 221 *min*, respectively. Similar execution times without significant performance loss were observed when 4 concurrent simulations were executed in the same compute node (scenario 1N4S), with average total times around 22 and 210 *min* for the *coarse* and *fine* meshes, respectively. A similar behaviour was observed using the MPI batch feature for the 128N512 scenario, with execution times of approximately 22 and 212 *min* for *coarse* and *fine* resolutions, respectively. Considering the times for the MPI process to start and end, these values were around 23 and 302 *min*, respectively. From the results in Table 1, it also transpires that the most demanding component is the solution of the ODE system of the myocardium, contributing around 11 and 94 *min* for the *coarse* and *fine* mesh resolutions, respectively. This is explained due to the larger number of control volumes of this domain compared to the Purkinje one, making this section of the problem more computationally demanding and the overall bottleneck.

**Table 1.** Execution times (minutes) for the cardiac digital twin test for the 3 submission scenarios. *Total*: total time; *Write*: writing time; *ECG*: ECG computation time; *M*_*ODE*_ /*M*_*PDE*_ : time to solve the myocardium ODE/PDE system; *P*_*ODE*_ /*P*_*PDE*_ : time to solve the Purkinje ODE/PDE system; *MPI*_*end*_: time for the MPI process to finish.

| Mesh | Scenario | Total | Write | ECG | $M_{ODE}$ | $M_{PDE}$ | $P_{ODE}$ | $P_{PDE}$ | $MPI_{end}$ |
| --- | --- | --- | --- | --- | --- | --- | --- | --- | --- |
| Coarse | 1N1S | **15.15** | 0.08 | 0.07 | 7.78 | 4.27 | 0.20 | 1.01 | - |
| | 1N4S | **22.24 ± 0.21** | 0.07 ± 0.00 | 0.11 ± 0.00 | 10.94 ± 0.11 | 7.23 ± 0.09 | 0.23 ± 0.00 | 1.05 ± 0.01 | 23.15 |
| | 128N512S | **22.41 ± 0.32** | 0.07 ± 0.00 | 0.11 ± 0.02 | 11.06 ± 0.17 | 7.24 ± 0.15 | 0.23 ± 0.01 | 1.06 ± 0.01 | 23.55 |
| Fine | 1N1S | **221.49** | 0.52 | 0.42 | 97.62 | 87.70 | 0.26 | 1.13 | - |
| | 1N4S | **210.66 ± 0.32** | 0.50 ± 0.00 | 0.42 ± 0.00 | 93.80 ± 0.18 | 84.81 ± 0.05 | 0.39 ± 0.01 | 1.13 ± 0.00 | 213.13 |
| | 128N512S | **211.94 ± 7.33** | 0.49 ± 0.01 | 0.42 ± 0.00 | 93.87 ± 0.50 | 85.89 ± 5.56 | 0.37 ± 0.03 | 1.13 ± 0.01 | 302.29 |

Based on these results, the novel MPI batch feature illustrates the scalability and efficiency of the proposed cardiac solver to conduct human-based cardiac digital twin studies under cluster GPU environments, enabling the correct adjustment of sensitive Purkinje coupling parameters to physiological ranges for anterograde PMJ delays. Additional results on the performed cardiac digital twin simulations are presented in Supplementary Material section A.7. Of importance, they provide further evidence supporting the existence of multiple possible cardiac digital twins with a similar ECG (see Supplementary Figures S9 to S11 in section A.7, for example). These results also indicate that, while sustaining analogous ECGs, different combinations of Purkinje coupling parameters can generate distinct distributions of PMJ delays across the Purkinje terminals. Based on that, certain Purkinje terminals exhibit more variability in their associated PMJ delays, which might indicate a more important role of such PMJs to the whole ventricular activation. Another relevant finding from these simulations is the existence of an almost linear relation between the Purkinje coupling parameters and the appearance of propagation block, as analysed by Figures 3A, S7 (Supplementary Material section A.6) and S9A (Supplementary Material section A.7).

## Conclusion

In this work, we have presented an open source, high-performance GPU solver for electrophysiology simulations. By systematically evaluating the solver’s performance across various CPU/GPU configurations, we demonstrated its efficiency in solving the monodomain model under different scenarios. Specifically, our findings highlight that leveraging GPUs for both the non-linear system of ODEs and the parabolic PDEs significantly accelerates computations, particularly for complex human-based cellular models. These results contrast with our earlier study emphasizing space adaptivity, as we found that directly copying all required data structures to the GPU at the start of the simulation yields greater efficiency gains. Furthermore, the solver’s integration into a cardiac digital twin pipeline demonstrated its scalability in a GPU cluster environment. The ability to execute 512 simulations concurrently across 128 compute nodes, with execution times comparable to single-node setups, shows its robustness for large-scale studies. These simulations were instrumental in calibrating Purkinje coupling parameters to achieve physiological anterograde PMJ delays, showcasing the solver’s applicability in patient-specific modelling that consider the cardiac conduction system as an essential component of the model. Our results affirm the solver’s capability to perform large-scale monodomain simulations efficiently, including detailed modelling of the Purkinje-muscle-junctions, a feature that, to the best of our knowledge was not implemented in any other open source cardiac solver. Furthermore, the experiments presented in this work can be seamlessly expanded to a large cohort of virtual patients, which is a relevant step towards applicability in *in silico* clinical trials and therapy evaluation. This work provides a significant advancement in the field, offering an open source scalable tool for researchers to explore complex cardiac electrophysiology scenarios with remarkable efficiency.

## Data Availability

The open source cardiac solver is publicly available at https://github.com/rsachetto/MonoAlg3D_C. All the necessary configuration files, custom functions, post-processing scripts used during the current study, as well as, the biventricular mesh and Purkinje networks used for the cardiac digital twin application will be publicly available at a Zenodo repository to allow reproducibility once the manuscript has been accepted.

## Acknowledgements

This work was funded by a Wellcome Trust fellowship in Basic Biomedical Sciences to B.R. (214290/Z/18/Z), the EPSRC project CompBioMedX (EP/X019446/1), the CompBioMed2 Centre of Excellence in Computational Biomedicine grant agreements No. 675451 and No. 823712, R.S.O. acknowledges support from Fapemig grant No. APQ-00748-18 and UFSJ.

R.W.S. acknowledges support from Fapemig grant APQ-02445-24 and UFJF. A.B.O. acknowledges support from UK Research and Innovation grant No. 10110728. The U.S. Department of Energy’s (DOE) Innovative and Novel Computational Impact on Theory and Experiment (INCITE) Program awarded access to Polaris, under contract No. DE-AC02-06CH11357. For the purpose of open access, the authors have applied a Creative Commons Attribution (CC BY) public copyright licence to any Author Accepted Manuscript version arising from this submission.

## Author contributions statement

L.A.B.: Conceptualization, Methodology, Software, Investigation, Formal analysis, Writing − original draft, Writing − review & editing. R.S.O.: Conceptualization, Methodology, Software, Investigation, Formal analysis, Writing − original draft, Writing

− review & editing. J.C.: Software, Formal analysis, Writing − review & editing. Z.J.W.: Software, Formal analysis, Writing − review & editing. R.D.: Data curation, Formal analysis, Writing − review & editing. A.B.O.: Formal analysis, Writing − review & editing. R.W.S.: Conceptualization, Methodology, Investigation, Formal analysis, Writing − original draft, Writing − review & editing. B.R.: Conceptualization, Formal analysis, Funding acquisition, Resources, Writing − review & editing.

## Additional information

### Supplementary material

accompanies this paper.

## References

1. Jean-Quartier, C., Jeanquartier, F., Jurisica, I. & Holzinger, A. In silico cancer research towards 3R. BMC cancer 18, 1–12 (2018).

2. Delp, S. L. et al. OpenSim: open-source software to create and analyze dynamic simulations of movement. IEEE transactions on biomedical engineering 54, 1940–1950 (2007).

3. Musuamba, F. T. et al. Scientific and regulatory evaluation of mechanistic in silico drug and disease models in drug development: Building model credibility. CPT: Pharmacometrics & Syst. Pharmacol. 10, 804–825 (2021).

4. Fenton, F. H., Cherry, E. M., Hastings, H. M. & Evans, S. J. Multiple mechanisms of spiral wave breakup in a model of cardiac electrical activity. Chaos 12, 852–892 (2002).

5. Lopez-Perez, A., Sebastian, R. & Ferrero, J. M. Three-dimensional cardiac computational modelling: methods, features and applications. Biomed. engineering online 14, 1–31 (2015).

6. Dasí, A. et al. In-silico drug trials for precision medicine in atrial fibrillation: From ionic mechanisms to electrocardiogram-based predictions in structurally-healthy human atria. Front. Physiol. 13, 966046 (2022).

7. Oliveira, R. S. et al. Ectopic beats arise from micro-reentries near infarct regions in simulations of a patient-specific heart model. Sci. reports 8, 16392 (2018).

8. Li, L. et al. Towards enabling cardiac digital twins of myocardial infarction using deep computational models for inverse inference. IEEE Transactions on Med. Imaging (2024).

9. Corral-Acero, J. et al. The ‘Digital Twin’ to enable the vision of precision cardiology. Eur. Hear. J. 41, 4556–4564 (2020).

10. Niederer, S. A., Sacks, M. S., Girolami, M. & Willcox, K. Scaling digital twins from the artisanal to the industrial. Nat. Comput. Sci. 1, 313–320 (2021).

11. Viceconti, M. et al. In silico trials: Verification, validation and uncertainty quantification of predictive models used in the regulatory evaluation of biomedical products. Methods 185, 120–127 (2021).

12. Haissaguerre, M., Vigmond, E., Stuyvers, B., Hocini, M. & Bernus, O. Ventricular arrhythmias and the His-Purkinje system. Nat. Rev. Cardiol. 13, 155 (2016).

13. Kavanagh, P. Open source software: Implementation and management (Elsevier, 2004).

14. Vigmond, E., Dos Santos, R. W., Prassl, A., Deo, M. & Plank, G. Solvers for the cardiac bidomain equations. Prog. biophysics molecular biology 96, 3–18 (2008).

15. Vázquez, M. et al. Alya: Multiphysics engineering simulation toward exascale. J. computational science 14, 15–27 (2016).

16. Alnæs, M. et al. The FEniCS project version 1.5. Arch. numerical software 3 (2015).

17. Plank, G. et al. The openCARP simulation environment for cardiac electrophysiology. Comput. Methods Programs Biomed. 208, 106223 (2021).

18. Africa, P. C. et al. lifex-ep: a robust and efficient software for cardiac electrophysiology simulations. BMC bioinformatics 24, 389 (2023).

19. Sachetto Oliveira, R. et al. Performance evaluation of GPU parallelization, space-time adaptive algorithms, and their combination for simulating cardiac electrophysiology. Int. J. for Numer. Methods Biomed. Eng. 34, e2913 (2018).

20. Gouvêa de Barros, B., Sachetto Oliveira, R., Meira Jr., W., Lobosco, M. & Weber dos Santos, R. Simulations of complex and microscopic models of cardiac electrophysiology powered by Multi-GPU platforms. Comput. Math. Methods Medicine 2012, 824569 (2012).

21. Sachetto Oliveira, R. et al. Comparing CUDA, OpenCL and OpenGL implementations of the cardiac monodomain equations. In Wyrzykowski, R., Dongarra, J., Karczewski, K. & Wasńiewski, J. (eds.) Parallel Processing and Applied Mathematics, 111–120 (Springer Berlin Heidelberg, Berlin, Heidelberg, 2012).

22. Amorim, R. M. & Weber dos Santos, R. Solving the cardiac bidomain equations using graphics processing units. J. Comput. Sci. 4, 370–376 (2013).

23. Kaboudian, A., Cherry, E. M. & Fenton, F. H. Large-scale interactive numerical experiments of chaos, solitons and fractals in real time via GPU in a web browser. Chaos, Solitons & Fractals 121, 6–29 (2019).

24. Biasi, N., Seghetti, P., Parollo, M., Zucchelli, G. & Tognetti, A. A matlab toolbox for cardiac electrophysiology simulations on patient-specific geometries. Comput. Biol. Medicine 185, 109529 (2025).

25. Vergara, C. et al. A coupled 3D-1D numerical monodomain solver for cardiac electrical activation in the myocardium with detailed Purkinje network. J. Comput. Phys. 308, 218–238 (2016).

26. Sundnes, J. et al. On the computational complexity of the bidomain and the monodomain models of electrophysiology. Annals biomedical engineering 34, 1088–1097 (2006).

27. Mazumder, S. Numerical methods for partial differential equations: finite difference and finite volume methods (Academic Press, 2015).

28. Gomes, J. M. et al. Uniformization method for solving cardiac electrophysiology models based on the markov-chain formulation. IEEE Transactions on Biomed. Eng. 62, 600–608 (2015).

29. Zipes, D. P., Jalife, J. & Stevenson, W. G. Cardiac electrophysiology: from cell to bedside E-book (Elsevier Health Sciences, 2017).

30. Behradfar, E., Nygren, A. & Vigmond, E. J. The role of Purkinje-myocardial coupling during ventricular arrhythmia: a modeling study. PLoS One 9, e88000 (2014).

31. Wiedmann, R. T., Tan, R. C. & Joyner, R. W. Discontinuous conduction at Purkinje-ventricular muscle junction. Am. J. Physiol. Circ. Physiol. 271, H1507–H1516 (1996).

32. dos Santos, R. W. et al. ATX-II effects on the apparent location of M cells in a computational model of a human left ventricular wedge. J. cardiovascular electrophysiology 17, S86–S95 (2006).

33. Vigmond, E. J. & Clements, C. Construction of a computer model to investigate sawtooth effects in the Purkinje system. IEEE transactions on biomedical engineering 54, 389–399 (2007).

34. Bishop, M. J. & Plank, G. Bidomain ECG simulations using an augmented monodomain model for the cardiac source. IEEE transactions on biomedical engineering 58, 2297–2307 (2011).

35. Niederer, S. A. et al. Verification of cardiac tissue electrophysiology simulators using an N-version benchmark. Philos. Transactions Royal Soc. A 369, 4331–4351 (2011).

36. ten Tusscher, K. H., Noble, D. Noble, P.-J. & Panfilov, A. V. A model for human ventricular tissue. Am. J. Physiol. Circ. Physiol. 286, H1573–H1589 (2004).

37. Tomek, J. et al. Development, calibration, and validation of a novel human ventricular myocyte model in health, disease, and drug block. Elife 8, e48890 (2019).

38. Rush, S. & Larsen, H. A practical algorithm for solving dynamic membrane equations. IEEE Transactions on Biomed. Eng. 389–392 (1978).

39. Banerjee, A. et al. A completely automated pipeline for 3D reconstruction of human heart from 2D cine magnetic resonance slices. Philos. Transactions Royal Soc. A 379, 20200257 (2021).

40. Camps, J. et al. Inference of ventricular activation properties from non-invasive electrocardiography. Med. Image Analysis 73, 102143 (2021).

41. Camps, J. et al. Digital Twinning of the Human Ventricular Activation Sequence to Clinical 12-lead ECGs and Magnetic Resonance Imaging Using Realistic Purkinje Networks for in Silico Clinical Trials. Med. Image Analysis 94, 103108 (2024).

42. Camps, J. et al. Harnessing 12-lead ECG and MRI data to personalise repolarisation profiles in cardiac digital twin models for enhanced virtual drug testing. Med. Image Analysis 100, 103361 (2025).

43. Doste, R., Coppini, R. & Bueno-Orovio, A. Remodelling of potassium currents underlies arrhythmic action potential prolongation under beta-adrenergic stimulation in hypertrophic cardiomyopathy. J. Mol. Cell. Cardiol. 172, 120–131 (2022).

44. Trovato, C. et al. Human Purkinje in silico model enables mechanistic investigations into automaticity and pro-arrhythmic abnormalities. J. Mol. Cell. Cardiol. 142, 24–38 (2020).

45. Li, J., Zhang, H. & Boyett, M. Numerical analysis of conduction of the action potential across the Purkinje fibre-ventricular muscle junction. In Computing in Cardiology, 265–268 (2016).

46. Mendez, C., Mueller, W. J. & Urguiaga, X. Propagation of impulses across the Purkinje fiber-muscle junctions in the dog heart. Circ. research 26, 135–150 (1970).

47. Matsuda, K. Configuration of the transmembrane potential of the Purkinje-ventricular fiber junction and its analysis. Electrophysiol. Ultrastruct. Hear. 177–187 (1967).

