## Supplementary Material for "MonoAlg3D: Enabling Cardiac Electrophysiology Digital Twins with an Efficient Open Source Scalable Solver on GPU Clusters"

### A Supplementary Material

#### A.1 Finite volume method applied to the anisotropic monodomain model

In this section we describe how the FVM is applied to solve the anisotropic monodomain model for the myocardium. As explained in the main text, the reaction and diffusion terms of the model can be separated using operator splitting. This leads, per timestep, to the solution of the parabolic linear PDE:

$$\beta C_m \frac{\partial V_M}{\partial t} = \nabla \cdot (\sigma_{\mathbf{M}} \nabla V_M). \tag{1}$$

For the spatial discretisation of its diffusion term, we consider the relations:

$$J = -\sigma \nabla V, \tag{2}$$

where $J$ $(\mu A/cm^2)$ represents the density of intracellular current flow and

$$\nabla \cdot J = -I_v. \tag{3}$$

Here, $I_v$ $(\mu A/cm^3)$ is regarded a volumetric membrane current and corresponds to the right side of equation (1).

After defining the mesh geometry and partitioning the domain in control volumes, the FVM equations can be written. Integrating equation (3) over each discretised volume:

$$\int_{\Omega} \nabla \cdot J \, dv = -\int_{\Omega} I_v \, dv, \tag{4}$$

and applying the divergence theorem together with the original PDE for the given domain, it yields:

$$\beta C_m \int_{\Omega} \frac{\partial V}{\partial t} dv = -\int_{\Omega} I_v \, dv = \int_{\Omega} \nabla \cdot J \, dv = \int_{\partial\Omega} J \cdot \vec{n} \, ds, \tag{5}$$

where $\vec{n}$ represents the surface normal vector. This equation is the basic term for deriving the linear system of equations associated with the discretisation of the PDE problem.

Let us consider for simplicity a bidimensional uniform mesh, consisting of regular squares with a space discretisation $h_M$. Situated in the centre of each volume $(i, j)$ is a node, and the transmembrane potential $V_M$ is the variable of interest. Assuming that $I_v$ represents an averaged value in each square, and using the result from equation (10) from the main manuscript, we have:

$$\left( \beta C_m \frac{\partial V_M}{\partial t} \right) \Bigg|_{(i,j)} = \frac{-\int_{\partial\Omega_M} J_{i,j} \cdot \vec{n} \, ds}{h_M^2}. \tag{6}$$

The calculus of $J_{i,j}$ can be split as shown in Figure S1A as the sum of the flows on the four faces of the control volume:

$$\int_{\partial\Omega} J_{i,j} \cdot \vec{n} \, ds = (J_{x_{i+i/2,j}} - J_{x_{i-i/2,j}} + J_{y_{i,j+i/2}} - J_{y_{i,j-i/2}}) h_M, \tag{7}$$

where

$$\begin{bmatrix} J_x \\ J_y \end{bmatrix} = \begin{bmatrix} \sigma_x & \sigma_{xy} \\ \sigma_{xy} & \sigma_y \end{bmatrix} \cdot \begin{bmatrix} \frac{\partial V}{\partial x} \\ \frac{\partial V}{\partial y} \end{bmatrix}. \tag{8}$$

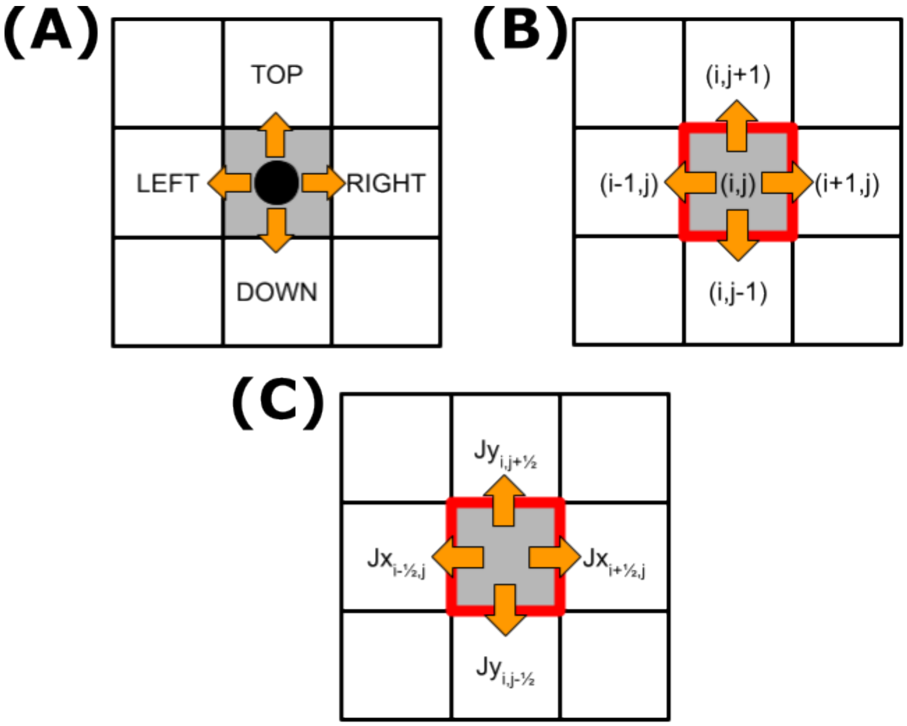


**Figure S1 .** Schematic of the 4 fluxes used to compute the intracellular current flow $J$ for a control volume in the bidimensional anisotropic case.

In terms of indexes, the mentioned flows passing through the faces of control volume $(i,j)$ are given by $(i,j+1)$, $(i,j-1)$, $(i+1,j)$ and $(i-1,j)$, respectively, as depicted in Figure S1B. In addition, the fluxes through each face are shown in Figure S1C and are given by:

$$\begin{aligned} J_{x_{i\pm1/2,j}} &= \sigma_{x_{i\pm1/2,j}} \left.\frac{\partial V}{\partial x}\right|_{i\pm1/2,j} + \sigma_{xy_{i\pm1/2,j}} \left.\frac{\partial V}{\partial y}\right|_{i\pm1/2,j}, \\ J_{y_{i,j\pm1/2}} &= \sigma_{xy_{i,j\pm1/2}} \left.\frac{\partial V}{\partial x}\right|_{i,j\pm1/2} + \sigma_{y_{i,j\pm1/2}} \left.\frac{\partial V}{\partial y}\right|_{i,j\pm1/2}. \end{aligned} \tag{9}$$

To calculate the flows in equation (9), the conductivity tensor needs to be evaluated at the faces of the control volume using the average value between the two neighbouring volumes:

$$\begin{aligned} \sigma_{x_{i\pm1/2,j}} &= \frac{\sigma_{x_{i\pm1,j}} + \sigma_{x_{i,j}}}{2}, \\ \sigma_{y_{i,j\pm1/2}} &= \frac{\sigma_{y_{i,j\pm1}} + \sigma_{y_{i,j}}}{2}, \\ \sigma_{xy_{i\pm1/2,j}} &= \frac{\sigma_{xy_{i\pm1,j}} + \sigma_{xy_{i,j}}}{2}, \\ \sigma_{xy_{i,j\pm1/2}} &= \frac{\sigma_{xy_{i,j\pm1}} + \sigma_{xy_{i,j}}}{2}, \end{aligned} \tag{10}$$

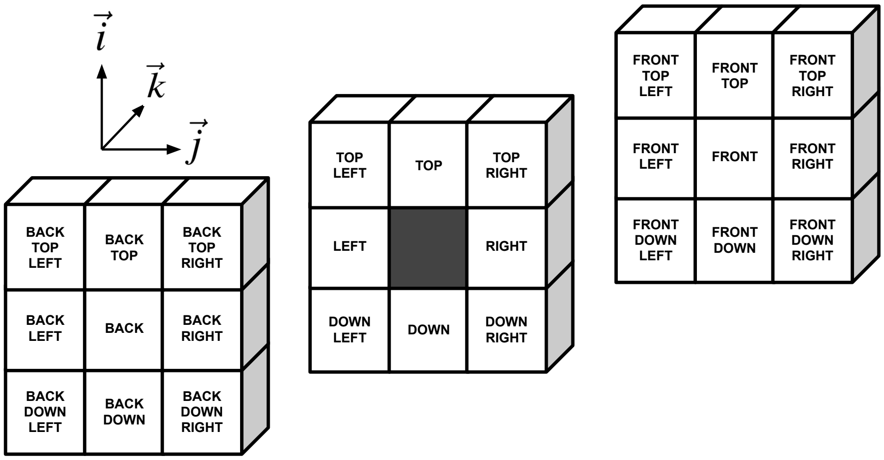


**Figure S2 .** Schematic of the 26 neighbourhood used to compute the intracellular current flow $J$ in the tridimensional anisotropic case.

while the spatial derivatives are approximated using a finite difference approach:

$$
\begin{aligned}
\left.\frac{\partial V}{\partial x}\right|_{i+1/2,j} &= \frac{(V_{i+1,j} - V_{i,j})}{h_M}, \\
\left.\frac{\partial V}{\partial x}\right|_{i-1/2,j} &= \frac{(V_{i,j} - V_{i-1,j})}{h_M}, \\
\left.\frac{\partial V}{\partial y}\right|_{i+1/2,j} &= \frac{(V_{i+1,j+1} + V_{i,j+1} - V_{i+1,j-1} - V_{i,j-1})}{4h_M}, \\
\left.\frac{\partial V}{\partial y}\right|_{i-1/2,j} &= \frac{(V_{i-1,j+1} + V_{i,j+1} - V_{i-1,j-1} - V_{i,j-1})}{4h_M}, \\
\left.\frac{\partial V}{\partial x}\right|_{i,j+1/2} &= \frac{(V_{i+1,j+1} + V_{i+1,j} - V_{i-1,j+1} - V_{i-1,j})}{4h_M}, \\
\left.\frac{\partial V}{\partial x}\right|_{i,j-1/2} &= \frac{(V_{i+1,j-1} + V_{i+1,j} - V_{i-1,j-1} - V_{i-1,j})}{4h_M}, \\
\left.\frac{\partial V}{\partial y}\right|_{i,j+1/2} &= \frac{(V_{i,j+1} - V_{i,j})}{h_M}, \\
\left.\frac{\partial V}{\partial y}\right|_{i,j-1/2} &= \frac{(V_{i,j} - V_{i,j-1})}{h_M}.
\end{aligned}
\tag{11}
$$

Similarly, for the tridimensional case, the equations follow an analogous pattern and can be derived using the same approach. The main differences are the inclusion in equation (7) of two additional flows in the $z$-axis ($J_{z_{i,j,k+1/2}}$ and $J_{z_{i,j,k-1/2}}$), and the application of a stencil considering all the 26 neighbouring control volumes, as shown in Figure S2.

#### A.2 Additional results for the cuboid benchmark

Figure S3 illustrates the propagation of excitation for the benchmark cuboid test (Figs. S3A-D), alongside corresponding action potentials from the *ten Tusscher* (Fig. S3E) and *ToR-ORd* (Fig. S3F) cellular models at the centre of the domain. Additionally, Figures S3G-H present the total number of control volumes to solve in the mesh over time when the space adaptivity was used for each model. As expected, when space adaptivity is active, the number of control volumes to be solved decreases as the domain starts to repolarise.

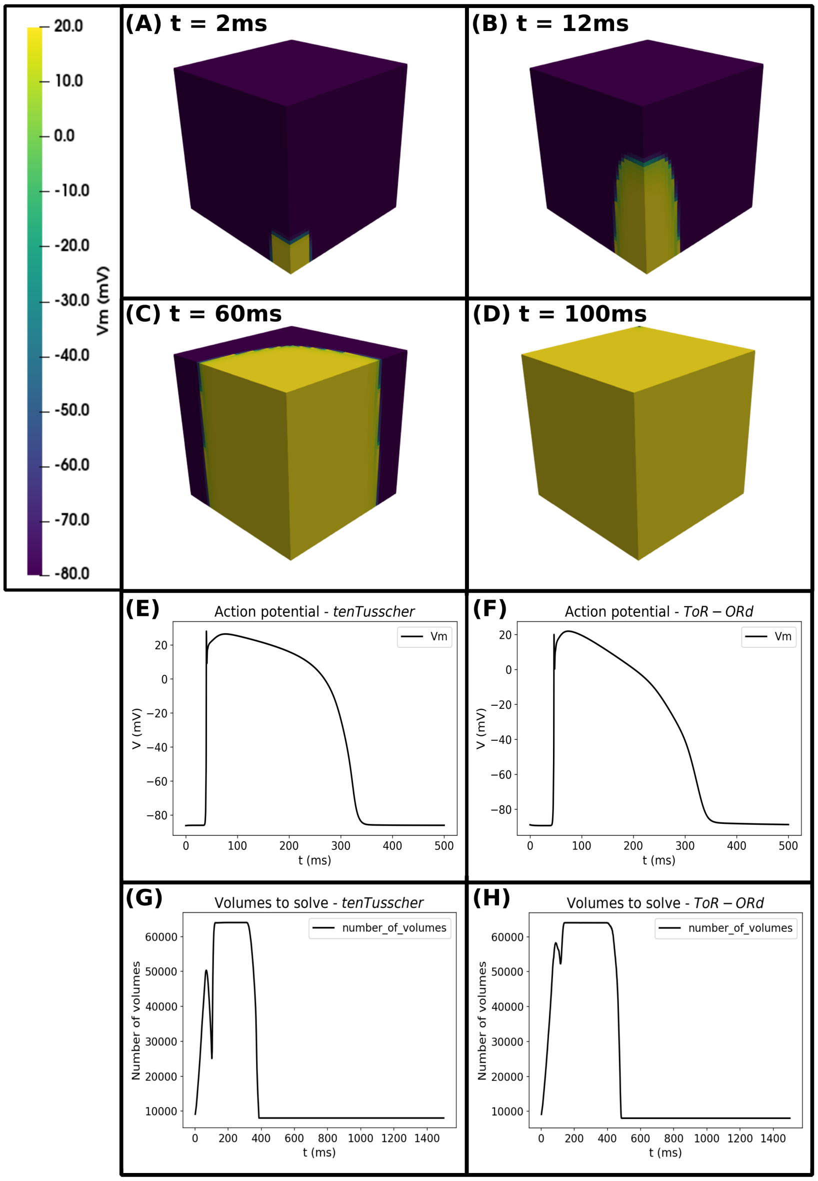


**Figure S3 .** Time evolution of transmembrane potential ($V_M$) for the benchmark cuboid test. (A-D) Snapshots at $t = 2$ $ms$, $t = 12$ $ms$, $t = 60$ $ms$, and $t = 100$ $ms$, respectively, when using the *ToR-ORd* cellular model. (E-F) Action potentials at the domain centre for the *ten Tusscher* and *ToR-ORd* cellular models, respectively. (G-H) Number of control volumes to solve in the mesh over time when using space adaptivity for both cellular models.

Tables S1 and S2 summarise the execution times for all the 6 scenarios and the two cellular models considered. Times spent on each portion of the solver are also detailed.

##### A.3 Cardiac digital twin mesh configuration

In these experiments, we use a human biventricular mesh (76 years old female, 87 $kg$, heart rate 48 $bpm$, 107 $cm^3$ volume), reconstructed from magnetic resonance imaging following[1]. This mesh has also been used in previous studies to personalise the subject's clinical ECG[2–4]. Further anatomical details are shown in Figure S4, including its coupling to the Purkinje network generated to activate the earliest activation sites inferred in[2,3] at given stimulation times (Fig. S4A), and the fibre orientation field of vector $\vec{f}$, equation (5) from the main manuscript (Fig. S4B). In addition, two thin layers are considered to model dense and sparse regions of subendocardial Purkinje coupling (Fig. S4C). Control volumes inside these regions are treated as isotropic (conductivity values calibrated based on the conduction velocities following[5], as detailed in Supplementary Material section

**Table S1 .** Execution times for the benchmark cuboid test in minutes (min) for the *ten Tusscher* cellular model in the 6 considered scenarios.

| Scenario | Total | Write | ODE | PDE | Reassemble matrix | Refine grid | Derefine grid | Order grid | Update cells | Update state vector |
|---|---|---|---|---|---|---|---|---|---|---|
| A+OC+PC | 5.82 | 0.42 | 2.42 | 1.99 | 0.13 | 0.38 | 0.32 | 0.10 | 0.01 | 0.00 |
| A+OG+PC | 4.40 | 0.42 | 0.48 | 2.00 | 0.13 | 0.40 | 0.34 | 0.10 | 0.01 | 0.35 |
| OC+PC | 9.51 | 0.89 | 7.21 | 1.36 | 0.00 | 0.00 | 0.00 | 0.00 | 0.00 | 0.00 |
| OC+PG | 8.56 | 0.88 | 7.18 | 0.41 | 0.00 | 0.00 | 0.00 | 0.00 | 0.00 | 0.00 |
| OG+PC | 2.91 | 0.88 | 0.62 | 1.33 | 0.00 | 0.00 | 0.00 | 0.00 | 0.00 | 0.00 |
| OG+PG | 1.77 | 0.85 | 0.42 | 0.39 | 0.00 | 0.00 | 0.00 | 0.00 | 0.00 | 0.00 |

**Table S2 .** Execution times for the benchmark cuboid test in minutes (min) for the *ToR-ORd* cellular model in the 6 considered scenarios.

| Scenario | Total | Write | ODE | PDE | Reassemble matrix | Refine grid | Derefine grid | Order grid | Update cells | Update state vector |
|---|---|---|---|---|---|---|---|---|---|---|
| A+OC+PC | 11.18 | 0.46 | 7.36 | 2.27 | 0.10 | 0.44 | 0.38 | 0.10 | 0.03 | 0.01 |
| A+OG+PC | 6.10 | 0.46 | 0.83 | 2.27 | 0.10 | 0.44 | 0.39 | 0.10 | 0.03 | 1.31 |
| OC+PC | 25.11 | 0.95 | 22.44 | 1.68 | 0.00 | 0.00 | 0.00 | 0.00 | 0.00 | 0.00 |
| OC+PG | 24.14 | 0.94 | 22.53 | 0.59 | 0.00 | 0.00 | 0.00 | 0.00 | 0.00 | 0.00 |
| OG+PC | 3.80 | 0.91 | 1.17 | 1.62 | 0.00 | 0.00 | 0.00 | 0.00 | 0.00 | 0.00 |
| OG+PG | 2.29 | 0.90 | 0.80 | 0.51 | 0.00 | 0.00 | 0.00 | 0.00 | 0.00 | 0.00 |

A.4), with a size for both layers equivalent to mesh discretisation element. Both regions are also used to generate a different number of PMJs on the endocardium for the Purkinje network generation method[6]. Finally, the scaling factor map for the $I_{Ks}$ current used in the myocardial cellular *ToR-ORd* model for T-wave personalisation[3] is shown in Figure S4D.

In terms of cellular electrophysiology, the above-mentioned *ToR-ORd* model was used for describing the myocardium, with minor modifications aiming to improve T-wave personalisation[4]: 50% scaling of $I_{Kr}$ conductance, 5-fold scaling of $I_{Ks}$ conductance[7], and reduction of the time constant of L-type calcium channel activation ($\tau_{jca}$) from 75 to 60 $ms$. For the Purkinje domain, the *Trovato* human Purkinje model[8] was considered. Both ODE systems were solved with a Rush-Larsen scheme and fixed timestep of $\Delta t = 0.01\ ms$. The same discretisation step was used to solve the associated PDEs, for a total simulation time of 600 $ms$. The stimulus protocol was a single pulse applied at the His bundle ($N_{cells} = 25$) of amplitude 40 $pA/pF$ and 2 $ms$ duration. Other monodomain parameters were set to $\beta = 1400\ cm^{-1}$ and $C_m = 1\ \mu F/cm^2$.

Next, to generate a full branched Purkinje network that sustains the morphology of the clinical ECG we used the open source Purkinje generation method as described in the work from Berg et al.[6]. The procedure consists of first generating additional PMJs within the dense and sparse regions of the endocardium, where these regions are highlighted in Figure S4C. Next, the new PMJs are connected by the extra branching procedure described in[6] using the minimum Purkinje network as the initial root.

The monodomain conductivities are calibrated to reproduce the conduction velocity (CV) and activation times given by the reduce-order model into a biophysically-detailed one (monodomain) using cable simulations with MonoAlg3D following the same CV tuning procedure as described in the work from Costa et al.[5].

Regarding the monodomain parameters utilized for this experiment, the surface-to-volume ratio was set to $\beta = 1400 cm^{-1}$ and the membrane capacitance to $Cm = 1 \mu F/cm^2$. The maximum simulation time is $t_{max} = 600 ms$ and a time discretization equal to $dt = 0.01 ms$ was used to solve the associated PDEs. For the ODEs system related to the Purkinje and myocardium cellular models, a Rush-Larsen scheme with a fixed timestep of $dt = 0.01 ms$ was applied.

To model the cellular dynamics of the myocardium domain, the adjusted version of the *ToR-ORd* human ventricular model[9] was used as previously described, while for the Purkinje domain, the *Trovato* human Purkinje model[8] was consider. The rationale to use both cellular models is that they can be considered the state-of-the-art in terms of human cellular model currently available in the literature.

#### A.4 Conductivities and conduction velocities

Conductivities were calibrated using monodomain cable simulations to match conduction velocities (CVs), as summarised in Table S3. Of note, all CVs are close to physiological values given by Durrer et al.[10]. The same CV tuning procedure presented in the work from Costa et al.[5] was applied with MonoAlg3D using the *tuneCV* script (https://github.com/rsachetto/MonoAlg3D_C/tree/master/scripts/tuneCV).

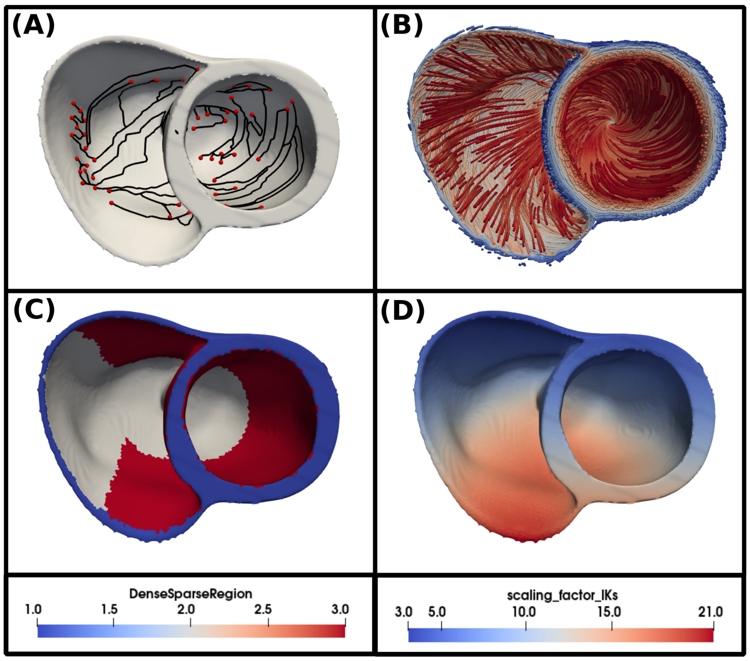


**Figure S4 .** Human-based biventricular mesh used for cardiac digital twin experiments. (A) Biventricular mesh coupled to the Purkinje network generated with the extra branching procedure from[6]. The Purkinje network is coloured in black, while red spheres illustrate the active PMJs. (B) Fibre orientation field of vector $\vec{f}$. (C) Heterogeneity in conductivity tensor $\sigma_M$, where blue denotes fully orthotropic conductivity values ($\sigma_f, \sigma_t, \sigma_n$), while white and red represent dense and sparse endocardial regions with isotropic conductivity values of $\sigma_{dense}$ and $\sigma_{sparse}$, respectively. (D) Scaling factor map of $I_{Ks}$ current in the *ToR-ORd* model used for T-wave personalisation[4].

**Table S3 .** Monodomain conductivity values ($mS/cm$) and conduction velocities ($m/s$) for the Purkinje and myocardium domains.

| | Purkinje | | Endocardium layers | | | | Fibre | | Sheet | | Normal | |
|---|---|---|---|---|---|---|---|---|---|---|---|---|
| | $CV_p$ | $\sigma_p$ | $CV_{dense}$ | $\sigma_{dense}$ | $CV_{sparse}$ | $\sigma_{sparse}$ | $CV_f$ | $\sigma_f$ | $CV_s$ | $\sigma_t$ | $CV_n$ | $\sigma_n$ |
| Coarse mesh | 3 | 115 | 1.3 | 8.81 | 0.97 | 5.35 | 0.65 | 2.83 | 0.27 | 0.92 | 0.48 | 1.83 |
| Fine mesh | 3 | 60 | 1.3 | 7.78 | 0.97 | 4.51 | 0.65 | 2.21 | 0.27 | 0.54 | 0.48 | 1.31 |

##### A.5 Cardiac digital twin simulation dispatch strategy

To dispatch the cardiac digital twin simulations on Polaris, we considered that optimal performance for simulations using entirely the GPU was attained with 8 OpenMP threads and 1 GPU device, as informed by the results from the benchmark cuboid experiments in Figure 2 from the main manuscript. (scenario OG+PG).

According to the hardware specification, a Polaris compute node has 32 physical CPU cores and 4 GPU devices. Within this context, we can execute 4 concurrent cardiac digital twin simulations on each node without wasting resources. To enable the 512 simulations, we made a job request for 128 compute nodes, where 4 jobs are concurrently executed on each node using the MPI batch feature (see Figure S5).

##### A.6 Purkinje-Muscle-Junction calibration

To validate the Purkinje module and enable a physiological range for its coupling parameters, a calibration experiment was conducted using two ventricular wedges from the considered biventricular mesh. Both wedges are activated by a single Purkinje terminal and are located at different regions of the left ventricle (Fig. S6A), in order to analyse the effects of the isotropic endocardial conductivities ($\sigma_{dense}$ and $\sigma_{sparse}$) on the anterograde PMJ delay. As so, the first wedge is located within the sparse endocardial region (Fig. S6B), while the second is in the dense endocardial region (Fig. S6C). Different space discretisations were also tested for both the Purkinje (100 and 250 $\mu m$) and the myocardial (250 and 500 $\mu m$) domains, with conductivities informed by our calibration results (Table S3). For the Purkinje coupling parameters, we considered a range of 25 equispaced

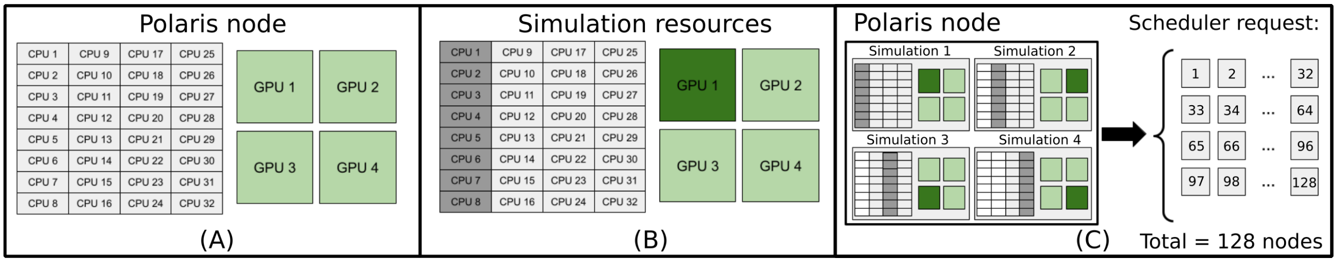


**Figure S5 .** Strategy to dispatch the cardiac digital twin jobs on Polaris. (A) Polaris node CPU/GPU hardware overview. (B) Amount of resources used for one cardiac digital twin simulation on Polaris. (C) Job allocation request to the machine scheduler to run 512 simulations using 128 nodes with the MPI feature.

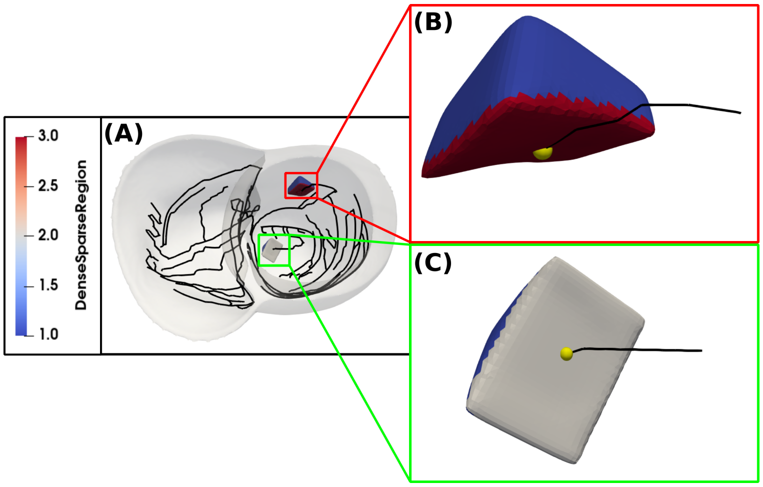


**Figure S6 .** Calibration of Purkinje coupling parameters $R_{PMJ}$ and $N_{PMJ}$. (A) Biventricular mesh with Purkinje network coloured in black, and the two considered ventricular wedges located in the sparse (red) and dense (grey) endocardial regions. (B) Close-up view of the ventricular wedge located in the sparse endocardial region. (C) Close-up view of the ventricular wedge located in the dense endocardial region.

values in $[100, 2500]$ $k\Omega$ for the PMJ resistance, $R_{PMJ}$, and 10 equispaced values in $[10, 100]$ for the number of myocardium control volumes linked to a terminal Purkinje control volume, $N_{PMJ}$.

Altogether, we performed $2,000$ simulations considering all possible combinations of wedges, space discretisations, and Purkinje coupling parameters. Simulations were executed for a total time of 50 $ms$, as we were only interested in measuring anterograde PMJ delays. These took $\approx 30$ $s$ to run enabled by the OG+PG setup. The PMJ delay was calculated as the time difference between the terminal Purkinje control volume and the closest myocardium control volume linked to it reaching a transmembrane potential threshold of $-40$ $mV$.

Figure S7 summarises the results for the anterograde PMJ delay, which is affected by different factors. For instance, the myocardial space discretisation decreases the anterograde PMJ delay as the mesh is refined. This is justified due to the fact that, for fixed Purkinje coupling parameters still activating the same number of myocardium control volumes, the size of the PMJ site will reduce for a finer myocardial mesh (less sink). This can be verified by pairwise comparisons of Figures S7A–S7C, S7B–S7D, S7E–S7G, and S7F–S7H.

Second, the Purkinje space discretisation also affects the anterograde PMJ delay. In this case, refining the Purkinje space resolution would increase the anterograde PMJ delay (less source). Equivalently, increasing the size of Purkinje control volumes increases the current flowing through the terminal volume face associated with the coupling, shortening the time necessary for the coupled myocardial volumes to depolarise. This can be visualised by comparing Figures S7A–S7E, S7B–S7F, S7C–S7G, and S7D–S7H.

Another parameter affecting the anterograde PMJ delay is the endocardial conductivity, with longer delays in the denser endocardial regions (Figs.S7A–S7B). This behaviour can be explained by the higher conductivity of the dense endocardium layer, leading to a larger diffusion of the current entering the myocardium through the Purkinje terminals to their neighbours, and hence an increase in the time needed for the myocardial cells to reach their activation threshold. The same conclusion can

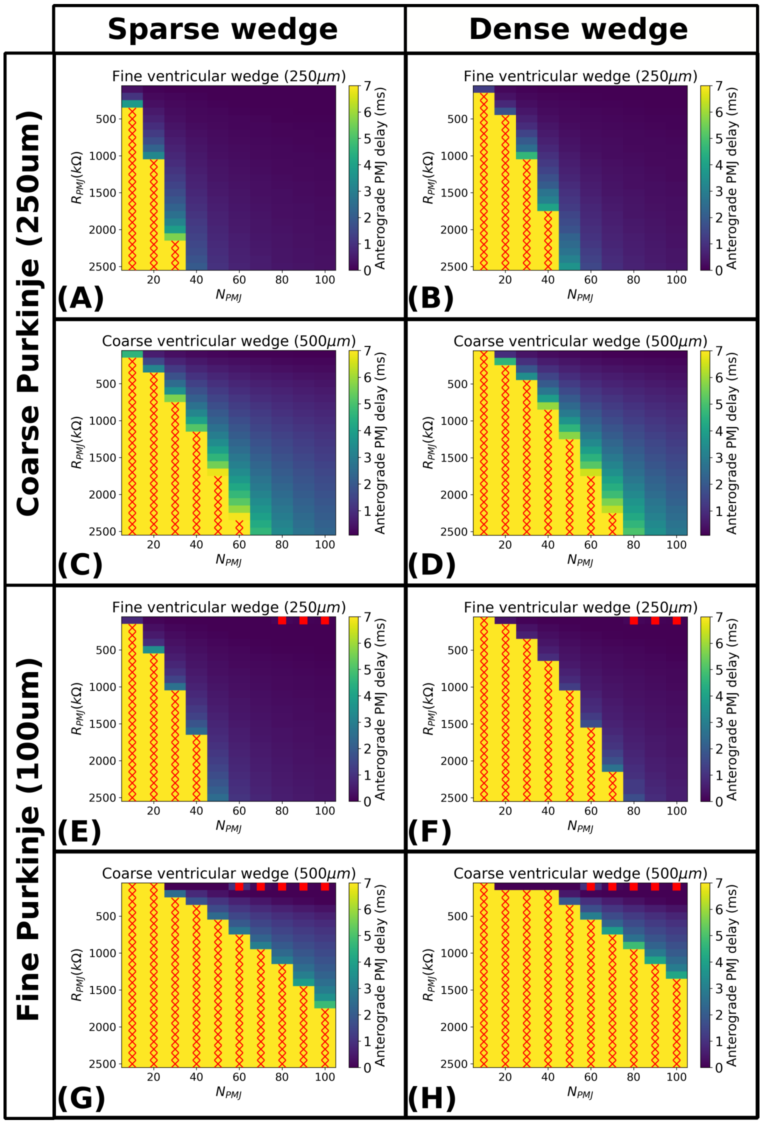


**Figure S7 .** Heatmaps of anterograde PMJ delays for varying Purkinje coupling parameters $R_{PMJ}$ and $N_{PMJ}$ at different space discretisations. Red crosses denote propagation block at the PMJ site, while red squares denote instabilities associated to solving the Purkinje coupling explicitly. (A) Sparse endocardial region at fine discretisation, coupled to a coarse Purkinje fibre. (B) Dense endocardial region at fine discretisation, coupled to a coarse Purkinje fibre. (C) Sparse endocardial region at coarse discretisation, coupled to a coarse Purkinje fibre. (D) Dense endocardial region at coarse discretisation, coupled to a coarse Purkinje fibre. (E) Sparse endocardial region at fine discretisation, coupled to a fine Purkinje fibre. (F) Dense endocardial region at fine discretisation, coupled to a fine Purkinje fibre. (G) Sparse endocardial region at coarse discretisation, coupled to a fine Purkinje fibre. (H) Dense endocardial region at coarse discretisation, coupled to a fine Purkinje fibre. Fine spatial discretisation: $h_M = 250\ \mu m$, $h_P = 500\ \mu m$. Coarse spatial discretisation: $h_M = 500\ \mu m$, $h_P = 250\ \mu m$. Conductivity values as per Table S3.

be drawn by pairwise comparisons of Figures S7C–S7D, S7E–S7F, and S7G–S7H.

Regarding the Purkinje coupling parameters, a balance exists allowing to achieve similar anterograde PMJ delays under different combinations of $R_{PMJ}$ and $N_{PMJ}$. For instance, the anterograde PMJ delay is proportional to the resistance of the PMJ site, as increasing $R_{PMJ}$ not only increases the delay but can lead to propagation block (Fig.S7, red crosses). The reason behind this is that the larger the resistance, the lesser current will flow through the terminal Purkinje control volume to the coupled myocardium volumes, making ventricular depolarisation harder at the PMJ sites. Alternatively, increasing the number of coupled myocardium control volumes has the opposite effect, decreasing the anterograde PMJ delay as more myocardium control volumes are coupled to the terminal Purkinje volume. The main reason is that $N_{PMJ}$ effectively affects the size of the PMJ site, as a result of the Purkinje terminals stimulating more myocardium control volumes. The same current that

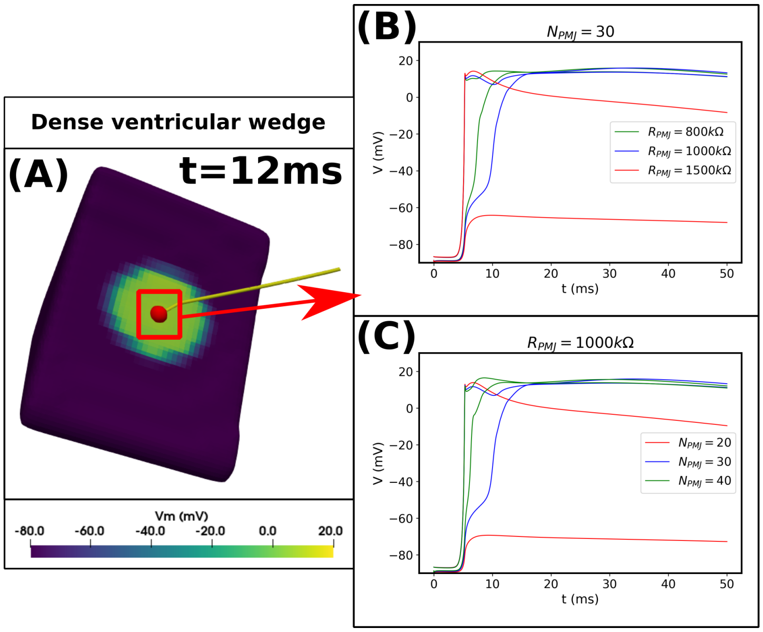


**Figure S8 .** Effect of the Purkinje coupling parameters on the transmembrane potentials of the Purkinje terminal and its closest coupled myocardium control volume. (A) Snapshot of the simulation for the wedge in the dense endocardial region, with $h_M = 250\ \mu m$, $h_P = 250\ \mu m$, $R_{PMJ} = 1000\ k\Omega$ and $N_{PMJ} = 30$. (B) Transmembrane potentials for fixed $N_{PMJ} = 30$ and varying $R_{PMJ}$, where the potentials of both the terminal Purkinje and myocardium control volumes are coloured in red for $R_{PMJ} = 1500\ k\Omega$, blue for $R_{PMJ} = 1000\ k\Omega$, and green for $R_{PMJ} = 800\ k\Omega$. (C) Transmembrane potentials for fixed $R_{PMJ} = 1000\ k\Omega$ and varying $N_{PMJ}$, where in red $N_{PMJ} = 20$, blue $N_{PMJ} = 30$, and green $N_{PMJ} = 40$.

flows through the Purkinje terminal control volume splits among the neighbouring myocardium control volumes. With more myocardium volumes being stimulated at the same time the depolarisation threshold can be reached through diffusion more quickly.

Further insights on impaired conduction at PMJ sites can be obtained by a deeper analysis of transmembrane potentials from the terminal Purkinje and its closest coupled myocardium control volume (Fig.S8). Keeping $N_{PMJ}$ fixed and increasing the resistance of the PMJ site (Fig.S8B), both the transmembrane potentials of the Purkinje and myocardial control volumes are affected by the electrotonic effects occurring at the domain interface. For instance, the wedge in the dense endocardial region is successfully stimulated for $R_{PMJ} = 1000\ k\Omega$ with an anterograde PMJ delay of approximately 5 $ms$. However, when the PMJ resistance is increased to $R_{PMJ} = 1500\ k\Omega$, the myocardium domain can not longer be properly stimulated by the Purkinje and the myocardium cells remain close to their resting state. A similar effect is observed when the number of coupled myocardium control volumes $N_{PMJ}$ is decreased while fixing the resistance of the PMJ (Fig.S8C). Such a behaviour on transmembrane potential has been observed experimentally in canine Purkinje fibre papillary muscle preparations, and its believed to be due to electrotonic interactions at PMJ junctions[11]. The stimulus through a PMJ site starts with an initial depolarisation of the Purkinje terminal, followed by a blunted upstroke. During this phase the Purkinje terminal is stimulating the coupled myocardial cells and raising their transmembrane potential due to the influx of current coming from the Purkinje fibres. The myocardial cells will depolarise if sufficient current exists to reach their activation threshold, but otherwise a propagation block arises.

##### A.7 Additional cardiac digital twin results

The results for the 512 biventricular simulation study using the *coarse* mesh setup are presented in Figures S9. These were used to select a pair of Purkinje coupling parameters $(R_{PMJ} = 1029\ k\Omega, N_{PMJ} = 43)$ providing both a physiological anterograde PMJ delay (Fig.S9A) and a sufficiently accurate approximation of the ECG (Fig.S9B). Figures S9D and S9F highlight the activation of a PMJ site exhibiting an anterograde delay of 5.61 $ms$ and a similar activation pattern as the one presented in Figure 5F from the main manuscript. Furthermore, on average, the anterograde PMJ delay was around $4.10 \pm 2.34$ $ms$, in agreement with the physiological range reported in the literature that is around $4 - 14$ $ms$[12]. Regarding the ECG approximation, we achieved an average PCC of 0.78 for this combination of parameters.

To further demonstrate how the Purkinje coupling parameters affect biventricular activation and the ECG, we selected 9 representative simulations from the 512 cases conducted for the *fine* mesh to analyse sensitivity to parameter changes. Figure

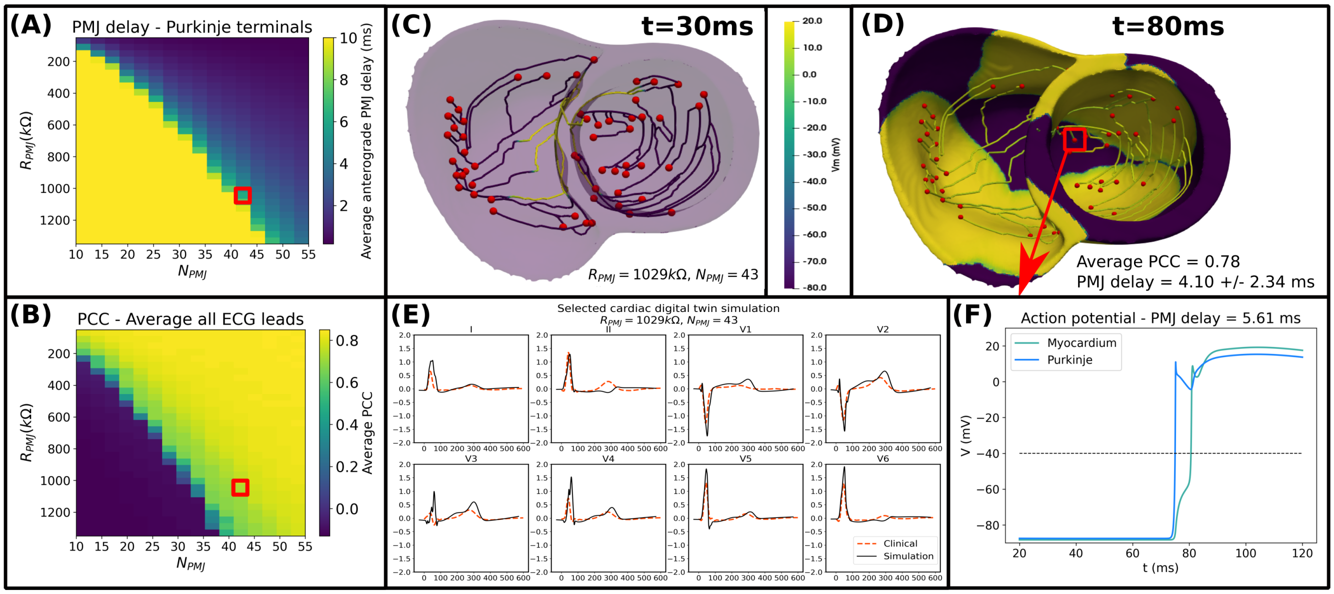

**Figure S9 .** Results for the 512 digital twin simulations with the *coarse* mesh. (A) Average anterograde PMJ delay across all Purkinje terminals. (B) Average PCC across all leads between the clinical and simulated ECG. Selected Purkinje coupling parameters ($R_{PMJ} = 1029\ k\Omega, N_{PMJ} = 43$) are highlighted by red squares. (C-D) Selected cardiac digital simulation with an average PCC of 0.78 in ECG reconstruction and average anterograde PMJ delay of $4.10 \pm 2.34\ ms$, at times $t = 30$ and $t = 80\ ms$, respectively. (E) Comparison between clinical and simulated ECGs. (F) Action potential upstrokes for the PMJ site highlighted in panel (D) with anterograde PMJ delay of 5.61 $ms$, at the terminal Purkinje volume (blue) and its closest coupled myocardial volume (lime).

S10 exemplifies that, depending on the values of the Purkinje coupling parameters, we can achieve both normal and impaired propagation at different PMJ sites, in agreement with experimental studies in canine Purkinje fibres[11,13].

Another interesting result is presented in Figure S11A, which illustrates the ECGs from 11 cardiac digital twin simulations using the *fine* mesh, all with an average PCC of 0.81. All lead traces are remarkably similar to each other, with only slight perceptual differences on the precordial leads near the start of the QRS complex and around the T-wave peak. The analysis of local activation times (LATs) from the closest coupled myocardium control volume to each Purkinje terminal also reveals that, in almost all terminals, the average LAT does not significantly change (Fig. S11B). This is however with the exception of a small number of Purkinje terminals, which present larger LAT variability of up to 9.11 $ms$. Moreover, the analysis of PMJ delays revealed even a larger variability across Purkinje terminals (Fig. S11C), still sustaining a similar ECG. These results hence reinforce the idea of a range of possible cardiac digital twins for a given patient.

A further surprising detail that can be noticed in Figure S11C is the presence of retrograde propagation in three Purkinje terminals. Figure S12 illustrates, for a *fine* mesh simulation using $R_{PMJ} = 1777\ k\Omega$ and $N_{PMJ} = 38$, one of the Purkinje terminals that exhibits retrograde propagation. As it can be appreciated in Figures S12A–B, a depolarising wave coming from the myocardium stimulates the Purkinje terminal before it gets activated by the Purkinje fibre. The analysis of the transmembrane potentials of the Purkinje terminal and its closest myocardial control volume (Fig. S12C) further verified a characteristic PMJ delay on the retrograde direction of 4 $ms$, which is close to the physiological range of $2-4\ ms$ reported on the literature [12,14]. Furthermore, the activation wave that stimulates those Purkinje terminals exhibiting retrograde propagation comes from the depolarisation of a right ventricular region. This activation wave then proceeds transmurally to the left ventricle across the ventricular septum, reaching the highlighted area in Figure S12A. Nevertheless, we did not observe any form of macro-reentry for this single cycle.

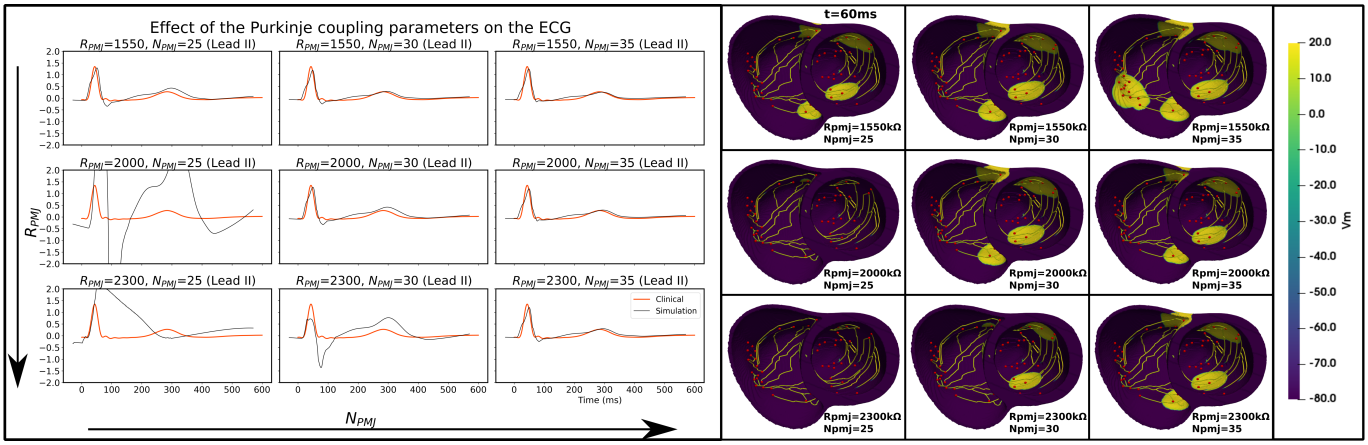


**Figure S10 .** Effects of Purkinje coupling parameters $R_{PMJ}$ and $N_{PMJ}$ on the simulated ECG (Lead II) and on biventricular activation, considering the *fine* mesh. The left panel illustrates 9 simulated ECGs (black traces) for different combinations of $R_{PMJ}$ and $N_{PMJ}$, compared against the clinical one (in red). The right panel illustrates the activation pattern at time $t = 60\ ms$ for each of the 9 corresponding simulations.

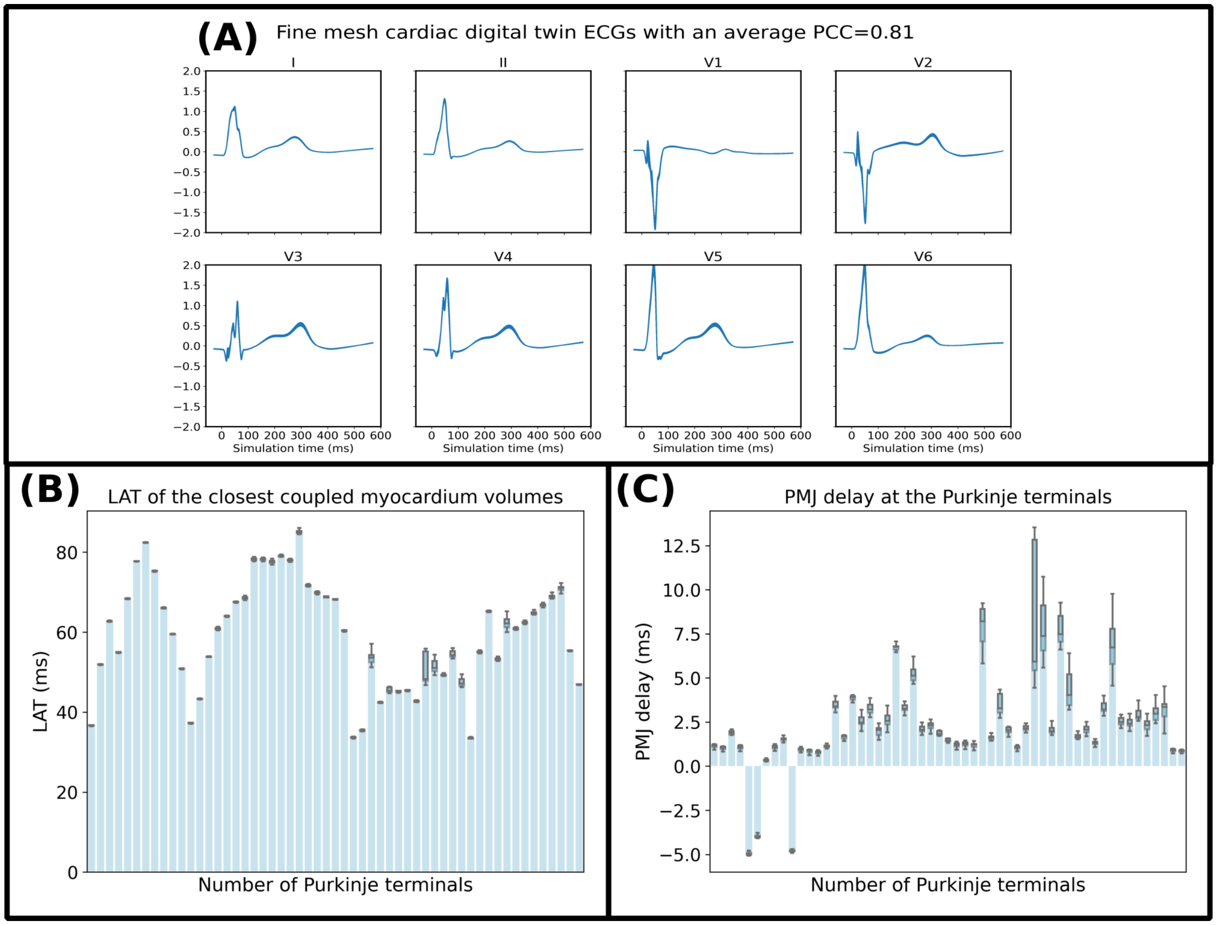


**Figure S11 .** (A) Comparison between 22 cardiac digital twin simulations using the *fine* mesh and average PCC of 0.81 for different combinations of the Purkinje coupling parameters. (B) Boxplots of local activation times $(ms)$ from the closest coupled myocardial control volumes to each of the 55 terminals located in the generated Purkinje network. (C) Boxplots of PMJ delay $(ms)$ for each Purkinje terminal.

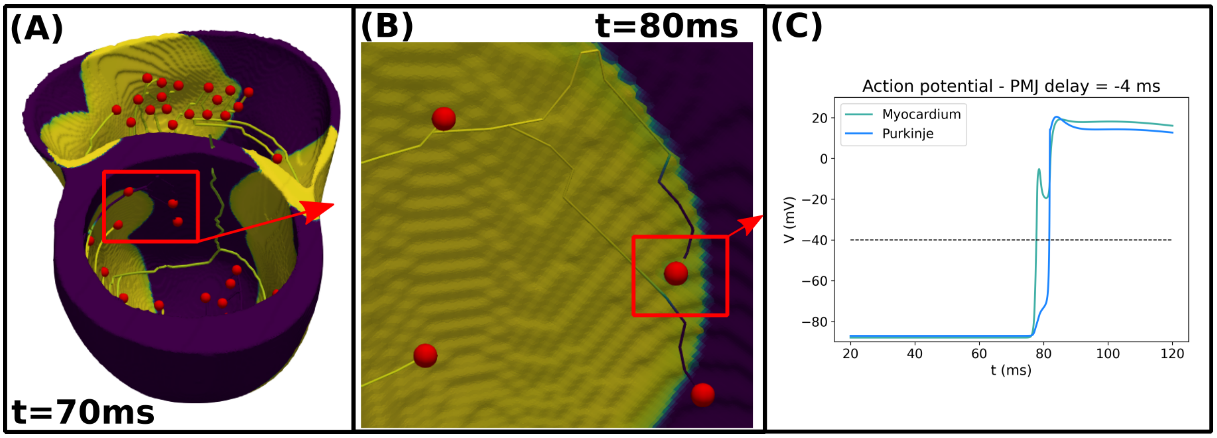


**Figure S12 .** (A) Region on the left ventricle where a Purkinje terminal is activated by retrograde propagation at time $t = 70$ *ms*. (B) Focused view of the Purkinje terminal activation at time $t = 80$ *ms*. (C) Transmembrane potential of the Purkinje terminal and closest myocardium control volumes from the highlighted PMJ site.

#### References


1. Banerjee, A. *et al.* A completely automated pipeline for 3D reconstruction of human heart from 2D cine magnetic resonance slices. *Philos. Transactions Royal Soc. A* **379**, 20200257 (2021).
2. Camps, J. *et al.* Inference of ventricular activation properties from non-invasive electrocardiography. *Med. Image Analysis* **73**, 102143 (2021).
3. Camps, J. *et al.* Digital Twinning of the Human Ventricular Activation Sequence to Clinical 12-lead ECGs and Magnetic Resonance Imaging Using Realistic Purkinje Networks for in Silico Clinical Trials. *Med. Image Analysis* **94**, 103108 (2024).
4. Camps, J. *et al.* Harnessing 12-lead ECG and MRI data to personalise repolarisation profiles in cardiac digital twin models for enhanced virtual drug testing. *Med. Image Analysis* **100**, 103361 (2025).
5. Costa, C. M., Hoetzl, E., Rocha, B. M., Prassl, A. J. & Plank, G. Automatic parameterization strategy for cardiac electrophysiology simulations. In *Computing in Cardiology 2013*, 373–376 (IEEE, 2013).
6. Berg, L. A. *et al.* Enhanced optimization-based method for the generation of patient-specific models of Purkinje networks. *Sci. Reports* **13**, 11788 (2023).
7. Doste, R., Coppini, R. & Bueno-Orovio, A. Remodelling of potassium currents underlies arrhythmic action potential prolongation under beta-adrenergic stimulation in hypertrophic cardiomyopathy. *J. Mol. Cell. Cardiol.* **172**, 120–131 (2022).
8. Trovato, C. *et al.* Human Purkinje in silico model enables mechanistic investigations into automaticity and pro-arrhythmic abnormalities. *J. Mol. Cell. Cardiol.* **142**, 24–38 (2020).
9. Tomek, J. *et al.* Development, calibration, and validation of a novel human ventricular myocyte model in health, disease, and drug block. *Elife* **8**, e48890 (2019).
10. Durrer, D. *et al.* Total excitation of the isolated human heart. *Circulation* **41**, 899–912 (1970).
11. Mendez, C., Mueller, W. J. & Urguiaga, X. Propagation of impulses across the Purkinje fiber-muscle junctions in the dog heart. *Circ. research* **26**, 135–150 (1970).
12. Behradfar, E., Nygren, A. & Vigmond, E. J. The role of Purkinje-myocardial coupling during ventricular arrhythmia: a modeling study. *PLoS One* **9**, e88000 (2014).
13. Mendez, C., Mueller, W. J., Merideth, J. & Moe, G. K. Interaction of transmembrane potentials in canine Purkinje fibers and at Purkinje fiber-muscle junctions. *Circ. research* **24**, 361–372 (1969).
14. Haissaguerre, M., Vigmond, E., Stuyvers, B., Hocini, M. & Bernus, O. Ventricular arrhythmias and the His-Purkinje system. *Nat. Rev. Cardiol.* **13**, 155 (2016).
